# Measuring effects of trainee professional development on research productivity: A cross-institutional meta-analysis

**DOI:** 10.1101/2020.09.28.316422

**Authors:** Patrick D. Brandt, Susi Sturzenegger Varvayanis, Tracey Baas, Amanda F. Bolgioni, Janet Alder, Kimberly A. Petrie, Isabel Dominguez, Abigail M. Brown, C. Abigail Stayart, Harinder Singh, Audra Van Wart, Christine S. Chow, Ambika Mathur, Barbara M. Schreiber, David A. Fruman, Brent Bowden, Chris E. Holmquist, Daniel Arneman, Joshua D. Hall, Linda E. Hyman, Kathleen L. Gould, Roger Chalkley, Patrick J. Brennwald, Rebekah L. Layton

**Author notes:** Patrick D. Brandt, Rebekah L. Layton, Patrick J. Brennwald, corresponding authors. (#current affiliation: University of Texas at San Antonio). (#Current affiliation: University of Texas at San Antonio). (#Current affiliation: Brown University). (#Current affiliation: University of Chicago). Denotes equal contributions (Patrick J. Brennwald, Rebekah L. Layton).

## Abstract

PhD-trained scientists are essential contributors to the workforce in diverse employment sectors that include academia, industry, government, and non-profit organizations. Hence, best practices for training the future biomedical workforce are of national concern. Complementing coursework and laboratory research training, many institutions now offer professional training that enables career exploration and develops a broad set of skills critical to various career paths. The National Institutes of Health funded academic institutions to design innovative programming to enable this professional development through a mechanism known as Broadening Experiences in Scientific Training (BEST). Programming at the BEST awardee institutions included career panels, skill-building workshops, job-searching workshops, site visits, and internships. An initial concern was since doctoral training is lengthy and requires focused attention on dissertation research, having students participate in additional complementary training activities might lengthen time to degree and hamper student research productivity. To address this concern, using time to degree and publication records as measures of efficiency and productivity, metrics were analyzed from ten BEST awardee institutions. Comparing doctoral students who participated to those who did not, results revealed that across these diverse academic institutions, there were no differences in time to degree or manuscript output. Furthermore, a few institutions even demonstrated a positive correlation between participation in career and professional development activities and productivity. Our findings suggest that doctoral students should be encouraged to participate in career and professional development opportunities to ensure their preparedness for a variety of diverse and important careers in the workforce.

**Significance Statement:** Our study is unique in that it compiled doctoral degree durations at ten different universities, recorded individual participation in career and professional development activities in terms of dosage, and tracked individual engagement in real-time rather than relying on surveys sent to trainees after graduation. Participation in career and professional development activities, including internships, did not decrease efficiency or productivity. Our findings suggest that doctoral students should be encouraged to participate in career and professional development opportunities to ensure their preparedness for a variety of diverse and important careers in the workforce.

## Introduction

Scientific doctoral education provides technical and cognitive-skill training and enables students to establish a positive sense of personal identity while building professional networks (1).

Importantly, doctoral training provides graduates with career value in the workforce as employers increasingly recognize that employees with PhDs have advanced knowledge and skills that can enhance the organization’s productivity and reputation (1).

Three decades ago, one in three biomedical doctoral students could have expected to join the academic tenure track; however, employment trends have since shifted (2–4). Both the National Institutes of Health (NIH) and the National Science Foundation (NSF) estimate that the current percentage of PhD scientists in tenured or tenure-track positions fell to fewer than one in four (3, 5, 6). This relatively lower percentage of PhD scientists transitioning to tenure-track academic positions is ascribed to several factors. First, the number of doctoral students graduating in the biomedical sciences in the United States has steadily risen, almost quadrupling over the past fifty years (3, 5, 6). Second, the growth in employment of biomedical doctoral graduates during this same time period has occurred almost entirely in industrial sectors, with comparatively little growth in employment in academic and government jobs (5–8). Third, graduates are preferentially choosing careers in research and research-related careers beyond academia, and this has only recently been widely recognized by the biomedical academic community (9).

### Experiential career training

Acknowledging that a broad range of careers are pursued by PhD graduates, many doctoral programs are being redesigned or supplemented to include experiential learning and skill development to prepare students for the biomedical workforce (10). Institutional efforts to supplement PhD training in preparation for varied career outcomes have been bolstered by funding opportunities from federal agencies, such as the NIH Broadening Experiences in Scientific Training (BEST) program, the NSF Research Traineeship program, and supplements to the NIGMS Training Grant programs (11–13). This “value-added” training for skills such as communication, working in teams and leadership (to name a few) are beneficial to those aspiring to either academic or non-academic positions (1).

Experiential learning opportunities, including internships, allow students to consider various career paths, and these additional professional and career development activities fill gaps in research training. These opportunities equip students with skills required in the workforce, expose graduate students to different workplaces, and make them more desirable as job candidates across career types (1, 2, 8, 14–16, 49). More recently, professional societies offer workshops on specialized professional development topics such as science policy and communication (17, 18), entrepreneurship and biotech careers (19), and provide other professional development programs (20). The national call for professional and career development underscores the value of PhD-trained scientists who demonstrate a variety of skills that transcend job sectors, find satisfying careers, and contribute to the workforce, both within and beyond academia. A call to action extends beyond the biomedical arena to include the physical and social sciences, as well as the arts and humanities, and is especially relevant in light of pandemic-centered disruption to the job market and accompanying economic turmoil.

### Efficiency and Productivity: Time to Degree and Publications

The overall length of doctoral training has long been an issue of concern. The NIH and other funding agencies, as well as policy makers, recommend exploring ways to embed career training into graduate education and postdoctoral training without increasing the time in training (5, 6).

Indeed, doctoral programs struggle to shorten the time to degree, prevent attrition, and guide doctoral students to meaningful careers after training (21). More than 85% of graduate deans surveyed in Canada, the United Kingdom, and the United States have taken steps to establish supervisor guidelines to help PhD students complete their programs in a timely fashion (22). Amidst the drive to shorten doctoral training periods, a persistent and understandable faculty concern is that to add such programming during training might take focus away from the laboratory and could potentially slow research progress, which might negatively impact grant funding, publication output, and time to degree (23). Nonetheless, data across universities show that time to degree for US students has remained relatively stable over the past fifteen years (5, 24).

Despite these concerns, many faculty recognize both the importance of career development to assist trainees and that their own knowledge in this area is lacking, such that supplemental programming is valuable (25). Moreover, initiatives that have promoted professional skills to complement scientific development have shown a benefit to graduate education and have not impacted time to degree or publication output, as highlighted by program evaluations (26–30). Initial data compiled from the baseline cohort of NIH BEST graduate trainees did not show a difference in average time in PhD programs over the first 3 years of data collection compared to average time before BEST implementation (8). To further test this hypothesis, a robust empirical comparison is needed to fully examine the effects of participation in professional development on time to degree and publications.

Hence, ten NIH BEST awardee institutions tested whether participation in career development activities affected time to degree as well as productivity (measured by published manuscripts) of doctoral students. BEST was an NIH grant program that funded seventeen institutions across the country to develop programming that could bridge the gap between research training and the job market, a transformative effort to catalyze career development change nationally (31). Our study is unique in that it compiled doctoral degree durations at ten different universities, recorded individual participation in career and professional development activities in terms of dosage, and tracked individual engagement in real-time rather than relying on surveys sent to trainees after graduation. Each of these ten BEST institutions developed distinctive program formats and structures. Data collected from these unique programs show that there was no difference in publication output or time to degree for doctoral students who participated, even quite actively, in career and professional development activities during their academic training.

## Materials and Methods

### Participants – Institutions, Programs, and Trainees

#### Institutions

Ten BEST institutions participated in this study. Participating institutions include (listed in alphabetical order): Boston University, Cornell University, Rutgers University, University of California, Irvine, University of Chicago, University of North Carolina at Chapel Hill, University of Rochester, Vanderbilt University, Virginia Tech, and Wayne State University. These institutions include public/private, city/rural, multiple/single-campus locations, and medical school/non-medical school settings. Institutions’ BEST programs supported populations ranging from 280-1000+ doctoral trainees, and 80-500+ postdoctoral trainees. Note that while some institutions include postdoctoral trainee participation in their BEST programs, all productivity data from these trainees were excluded from this study because they do not have a time to degree, making it more difficult to make comparisons. Details characterizing institutional profiles, BEST programs, and graduate departments/programs included in the study are provided for each institution in Supplemental File 1 (**S1**).

Across institutions, the departments, programs, and disciplines included in this study ranged from a single biomedical PhD program to programs serving all biological and biomedical programs; some of the institutions also include engineering, public health, or psychology disciplines (**S1)**. Common programs included Molecular Biology, Genetics, Biochemistry, Biomedical Sciences, Neuroscience, to name a few - as can be seen visualized with a weighted word cloud based on participating departmental and program names (**S1)**.

#### Ethics

In accordance with institutional review board (IRB) approvals as exempt (BU IRB#: H-33268; Rutgers IRB#: E15-050; Rochester IRB#: RSRB00055304; UNC IRB# 14-0544; Vanderbilt IRB# 190288; UCI IRB#: 2014-1502; VT IRB#: 13-711; WSU IRB: #094013B3E; the remaining institutional studies were approved via IRB Exemption Protocol ID#: 1412005184 through NIH OMB #0925-0718), participation was completely voluntary and informed consent was attained by affirming participation during program surveys. Students were giving the opportunity to opt out. All identifying data has been removed as per IRB requirements. Data is accessible via <u>Open Science Framework repository</u>.

#### Program Activities

Each BEST institution developed its own program to achieve proposed program-specific goals. Program activities ranged from single events to multi-part workshop series or coursework, as well as experiential learning activities, such as site visits, internships, and individual training sessions. One-off workshops were the most common activity each year for all of the programs (8). Institutions also deployed a wide range of activities differently, allowing trainees to participate through specific phases, by sector, by career interests, *ad hoc*, or some combination thereof. Most institutions included experiential learning opportunities with partners outside the university. Many programs offered opportunities at their university by partnering with various professional schools, core facilities, or support offices within their institution. Another focus was on incorporating mentorship and connecting trainees to alumni and professionals in broad areas of biomedical research. From these internal and external institutional connections, a majority of the BEST institutions allowed the possibility of internships, but it was not a requirement. The BEST institutions shared strategies, activities, and contacts among the BEST network of institutions during annual NIH BEST conferences, allowing programmatic offerings to evolve over time. A more complete description of the BEST institutions’ programming can be found in Supplemental File 1 (**S1)**.

#### Procedures

During the duration of BEST funding, institutions collected data about biomedical PhD trainee time to defense and level of participation in internships and BEST activities (*e.g*., career panels, skill-building workshops, job-searching workshops, site visits, internships). Data were submitted annually to NIH over a five-year period using common forms, standardized data-collection procedures, and compatible reporting methods to allow for cross-institutional comparison. Meetings to discuss evaluation of program design were held with all BEST consortium members, including a data summit to finalize common definitions and standardize BEST data collection methods (8; detailed collection methods including baseline data survey design and results included). Cross-institutional definitions for methods of instruction/delivery and agreements on common criteria for data were instrumental in developing data collection methods.

The most straightforward comparison between participants and non-participants in BEST career and professional development programming was measurement of binary outcome differences.

Hence, this was the most reliable effect-size measure to use, and was employed for meta-analytic comparisons. For binary comparisons using a t-test, the no participation group (control) was compared with any participation (e.g., medium plus high participation groups), giving a sense of effect size.

We were also interested in identifying potential dose-response effects based on level of participation. As each institution offered different events with variable length and scope, each was asked to define low participation and high participation levels independently (**S1**). Most institutions split their low and high dosage populations based on the observed median dosage level. These definitions were established so that the three groups could be compared with ANOVA analysis, giving a sense of any dose-response effect. This additional level of analysis yielded a more nuanced ability to evaluate participation effects and query for potential negative effects on productivity when there were high levels of participation. Nonetheless to retain the clarity of the control vs participant populations, all cross-institutional analyses were based upon bivariate comparisons.

For all binary analysis, with one exception, control groups were defined as non-participants; the exception was one program that did not have a true control group and hence divided participation in BEST events into an approximation of a control group (0-1 points) and a medium/high dose, rather than the null, low, high dose used by the remaining institutions. For consistency, the comparison groups for ANOVA are referred to as control, low, and high (control* is used to denote the approximated control group). Post-hoc analysis shows no difference when this institution’s data was excluded, hence we chose to include the data to be comprehensive.

Institutions also collected and reported publication outcomes. These data were independently gathered by each institution. Publication data were collected either by self-reported survey, manual PubMed queries or using the PubMed API using a Python script developed for this purpose and freely provided by Daniel Arneman and Joshua Hall (see Supplemental File 4 **S4;** 47). For those institutions that used the Python script, results were manually spot-checked for potential errors (**S4**), including overcounts for common names, legal name changes, nickname use, or advisor switching. In addition, extreme publication counts identified by the automated script (e.g., 0 publications or >5 publications) were manually rechecked by hand.

#### Analyses

Binary participant/non-participant comparisons were evaluated using independent sample t-tests, whereas dose-response tests were run using a one-way analysis of variance (ANOVA) with a three-level professional development dose variable (control, low, high). All comparisons were analyzed using Prism GraphPad (v8.4.0) software, which was also used to generate plots throughout the manuscript. All p values are reported to two significant figures.

The use of meta-analyses allows for extrapolation of an effect size and significance across different populations, multiple studies, or in our case different institutions and interventions. Hence, meta-analyses are preferred when comparing effects, especially when the variables of interest are measured differently across sites (e.g., hours, events, points). Although it is not uncommon to have fewer than optimal sample numbers (32), this situation is not ideal for use with meta-analyses. Therefore, meta-analyses were conducted only when a large enough set of institutions provided data (9-10 studies per meta-analysis). In some cases, not enough institutions were able to provide data to allow for meta-analysis (*i.e*., only a subset of institutions supporting internships). Meta-analyses were performed by entering effect sizes, p-values, and sample sizes for each institution’s data on that variable into Jamovi (v1.2.16) to produce an overall analysis of whether there was a significant effect across the population.

Primary predictors included the amount of professional development participation (binary or control/low/high dosage). Primary outcome variables of interest included productivity as measured by time to degree and publications (total and first-author). Finally, all outcome measures were tested against internship participation, the highest dose of professional development implemented across sites for the subset of institutions able to provide this data.

Power calculations verified whether our sample sizes were sufficient to detect a small effect size across each type of meta-analysis (33, 34). Post-hoc power analyses determined that >80% power was achieved for each meta-analysis, indicating that a sufficient number of subjects and studies were included. Meta-analytic power was calculated in accordance with recommendations by Harrer, Cuijpers, Furukawa, & Ebert (35). Meta analysis results and power calculations are grouped with other relevant analyses.

## Results

### 1. Participation & Efficiency: Time to Degree

#### Efficiency: Time to Degree versus Professional Development Participation

As BEST programs were implemented at each institution, some in the biomedical training community questioned whether participation in professional development programming would increase time to degree. Here, we tested this hypothesis using binary measurements (participants versus non-participants), as well as using a dose-response effect to determine whether higher levels of participation affect time to degree. The t-tests were conducted for bivariate analyses, ANOVAs for multiple groups, and multiple comparisons were only conducted if warranted by a significant omnibus finding.

One institution showed a statistically significant shorter time to degree for participants using either the binary or dose/level of analysis; the remaining institutions showed no significant difference in time to degree for participants in the binary condition or when accounting for level of participation (**Figure 1a, 1b)**

**FIGURE 1.**
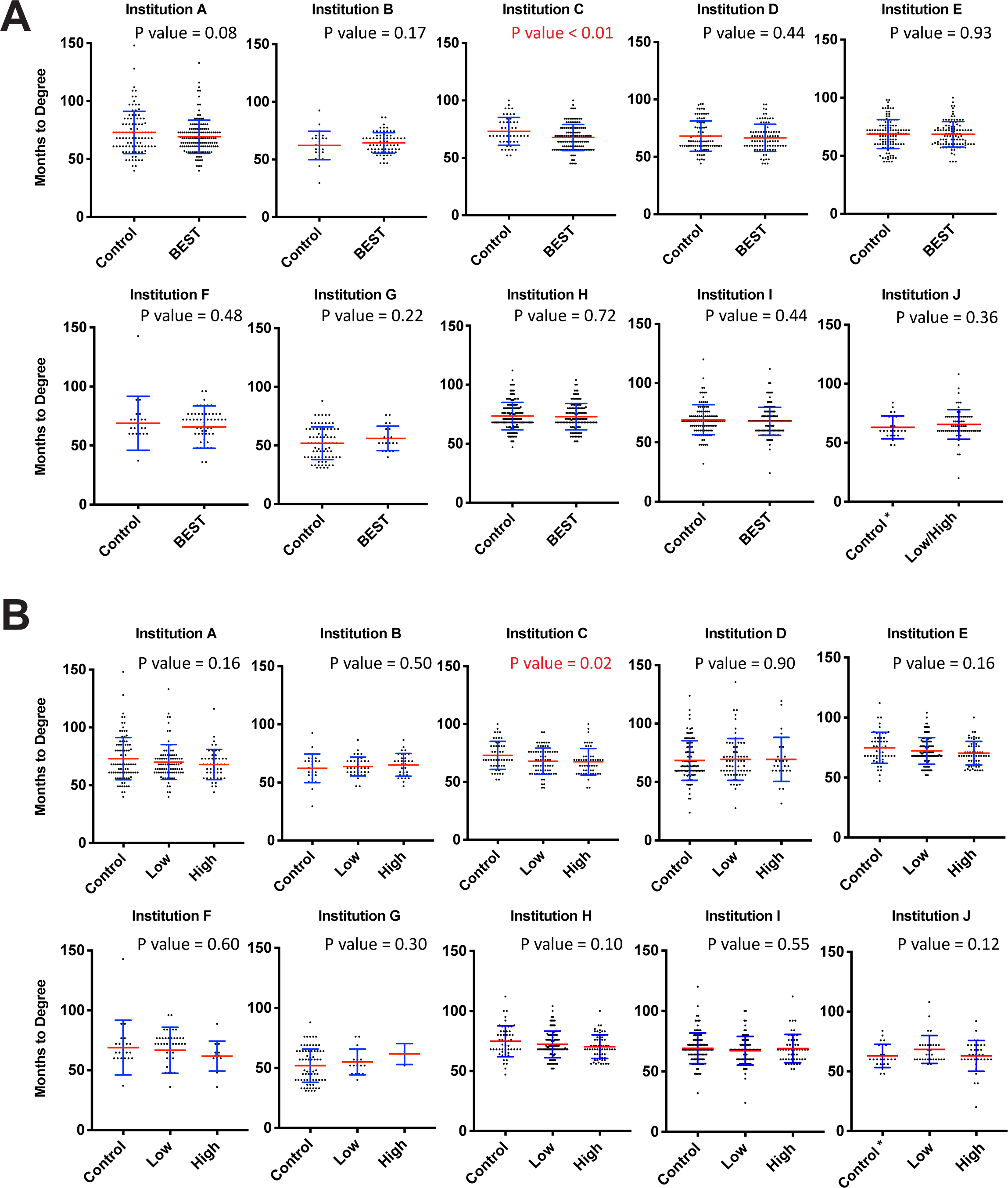
Professional Development Participation is Not Associated with an Increase in Time to Degree. A, Months to degree versus binary professional development participation. Blue error bars represent standard deviation of the mean. Mean is denoted by a red line. Independent samples t-tests were used to compare of control (non-participants) vs. participant time to degree (significant values of p<.05 noted in red). Control* for institution J indicates the control individuals were approximated based on available participation data (see Material and Methods). B, Months to degree versus dosage of professional development participation. Blue error bars represent standard deviation of the mean. Mean is denoted by a red line. Analysis of variance was used to compare the impact of control, low, and high dose participation on time to degree (significant values of p<.05 noted in red). Control* for institution J indicates the control individuals were approximated based on participation data (see Material and Methods). The remaining participants were divided into low and high participation groups.

Using the measure of months to defense resulted in one additional institution (*i.e.,* two total) showing that greater participation was associated with a statistically significant decrease in time to defense (**SI Figure 2a**). Overall, the data failed to support the hypothesis that participation in career and professional development at any level tested leads to a statistically significant increase in time in graduate training.

**Figure 2.**
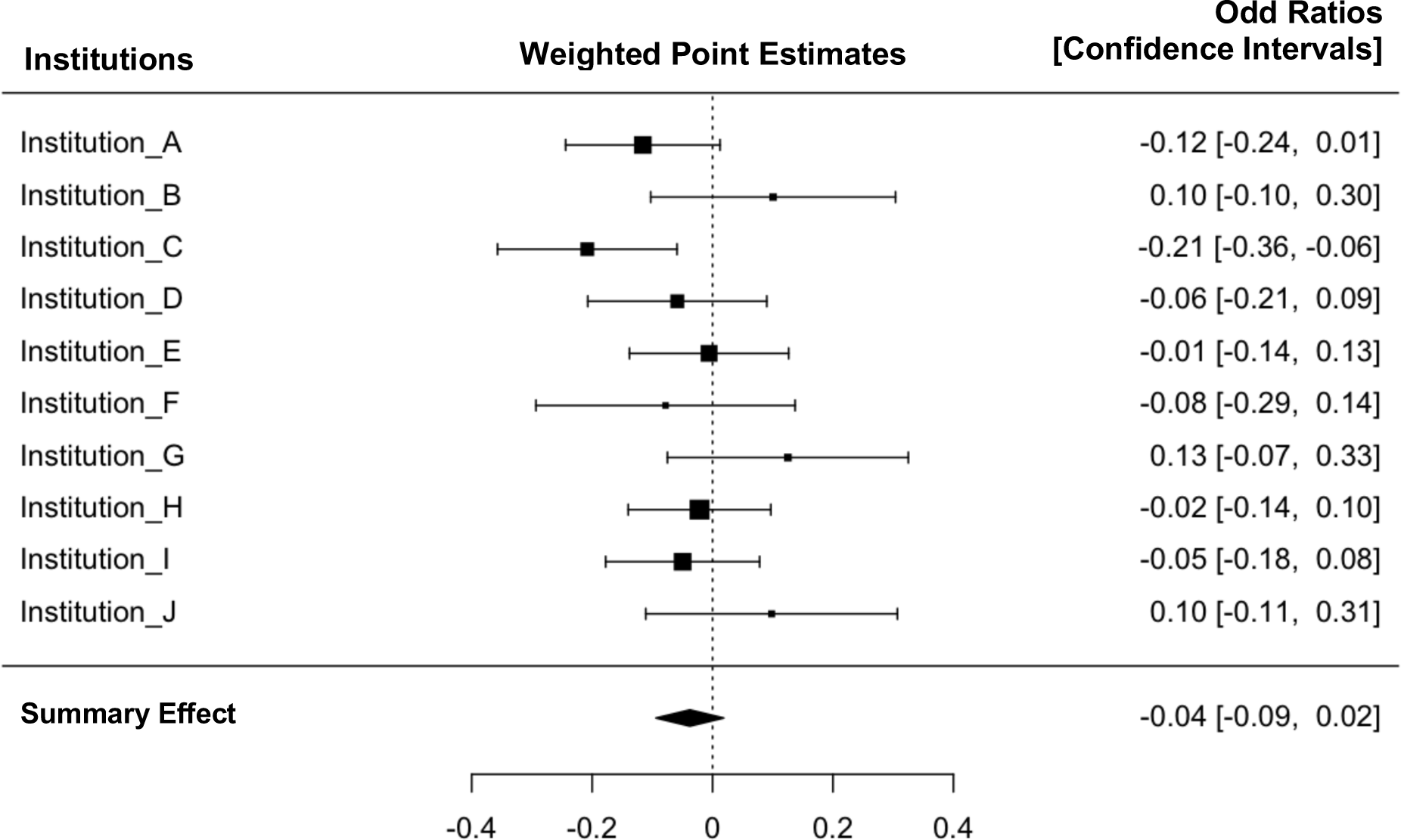
Graduate student efficiency measured by time to degree versus bivariate participation. Meta-analysis Forest plot displaying mean effect sizes (squares) and confidence intervals (brackets) for effect sizes of time to degree versus bivariate professional development participation (control versus participants). Large squares denote greater impact on the summary effect based on sample size and effect size in each institutional sample. The vertical dotted line represents a null effect. The size and shape of the diamond at the bottom of the Forest plot represents the summary effect. Because the diamond overlaps the vertical line (null effect), this indicates that the effect of professional development participation on time to degree is not significant.

#### Meta-Analysis of Effects on Trainee Efficiency

A meta-analysis (**Figure 2**) was conducted to determine a weighted effect size and significance across all the institutions. This cross site meta-analysis (including 1700 trainees’ participation data) showed no difference in time to degree between participants and non-participants, a point-estimate of -0.04, *p*=0.19, SE=0.03, z=-1.31, [CI95%= -0.09, 0.02], with effect sizes ranging from *r*^2^ = 0.01 - 0.04 (*r*s from -0.21 to +0.12). Power calculations suggest that with the average sample sizes and number of participating institutions’ data available for this study (N1=78, N2=95; alpha=0.05, k=10, *d*=0.20), and an observed low heterogeneity (I^2^ = 24.72% < 25% cutoff for low heterogeneity; τ^2^ =0.002, SEτ=0.004; Q=12.91, *p*=0.17) in a random effects model (35, 36), we had nearly 90% power to detect a small effect size (89%). Given that our study was cross-institutional and well exceeded the acceptable rate of 80% power (with an alpha of 0.05; 33, 34), we can confidently say that we had the ability to detect an effect size of this magnitude or greater.

Furthermore, there were no cases in which the dose-response effects were significantly longer for those with the highest participation (omnibus F-tests were not significant); in fact, in the single case of significant difference, the directionality indicated a favorable association such that participants took less time to graduate than non-participants. ANOVAs show comparisons between no-dose, low-dose, and high-dose event participation (**Figure 1b**).

In sum, the analysis reveals that participating in career and professional development was not associated with an increased time to degree. This initial finding supports the notion that participation, even in high doses, is not associated with any delay.

### 2. Participation and Productivity: Total Publications

Next we evaluated the impact of career and professional development participation on productivity, measured by number of publications. We first evaluated total publications during the graduate training period. For participants versus non-participants, one institution showed significantly more publications for participants, and one showed significantly fewer publications for participants. The remaining seven institutions showed no significant difference between participants and non-participants with regard to total number of publications, and when accounting for different levels of participation, no institution showed any significant difference in the number of total publications between groups (**Figure 3**).

**FIGURE 3.**
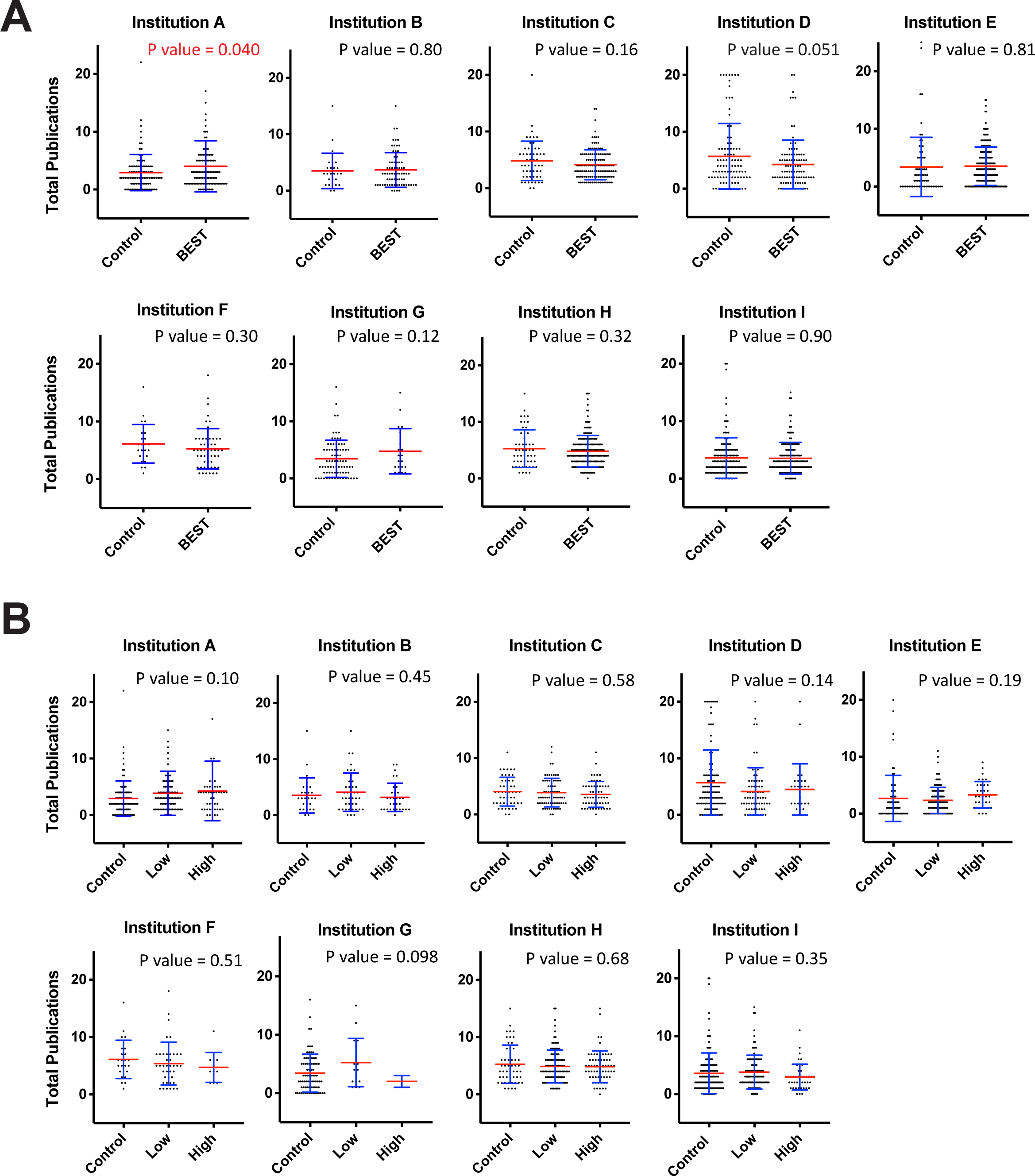
Professional Development Participation is Not Associated with a Decrease in Total Publication. A, Total publications versus binary professional development participation. Blue error bars represent standard deviation of the mean. Mean is denoted by a red line. Independent samples t-tests were used to compare of control vs. participant total publications (significant values of p<.05 noted in red). B, Total publications versus dosage of professional development participation. Blue error bars represent standard deviation of the mean. Mean is denoted by a red line. Analysis of variance was used to compare the impact of control, low, and high dose participation on total publications (significant values of p<.05 noted in red).

### 3. Participation & Productivity: First-author Publications

Professional scientists, faculty researchers, and doctoral training programs often place special significance on first-author publications because the bulk of trainees’ efforts in the lab are usually directed at projects resulting in first-author publications. These efforts also typically form the underpinning for the students’ theses. Due to the unique importance of first-author publications, we further examined whether there is a specific impact of participation in career and professional development on first-author publications.

Similar to the overall number of publications, there was no conclusive effect of BEST participation on increases or decreases in, specifically, first-author publications (**Figure 5**). In the binary condition for first-author publications, one institution’s BEST participants produced significantly fewer first-author publications. In contrast, when level of participation was considered, one institution’s “high dose” BEST participants produced significantly more first-author publications. In both the binary and dose-response analyses, the remaining eight institutions showed no significant difference between participants and non-participants in first-author publications. Accordingly, there was no overall trend of BEST participation reducing first-author publications, and the hypothesis that participation in professional development activities reduces publication rate was not supported by our data.

Meta-analyses were conducted to determine the weighted effect size and significance across all the institutions for total and first author publications (**Figures 4 and 6**). The cross-site meta-analyses (including nearly 1500 trainees’ publication data) showed no significant difference in total publications between participants and non-participants, with a point estimate of -0.04 (*p*=0.23, SE=0.03, *z*=-1.21, [CI95%= -0.10, 0.02]), with effect sizes ranging from r^2^ = < 0.01 -0.02 (**Figure 4**). Similarly, a meta-analysis of first-author publications from the same institutions showed no significant difference in first-author publications between participants, and non-participants, with a point-estimate of -0.02 (p=0.64, SE=0.04, z=-0.47, [CI95%= -0.08, 0.05]), with effect sizes ranging from r^2^ = < 0.01 - 0.03 (**Figure 6**). Across a large multi-institutional sample, collectively there was a lack of evidence for reduced trainee productivity as measured by publication number.

**Figure 4.**
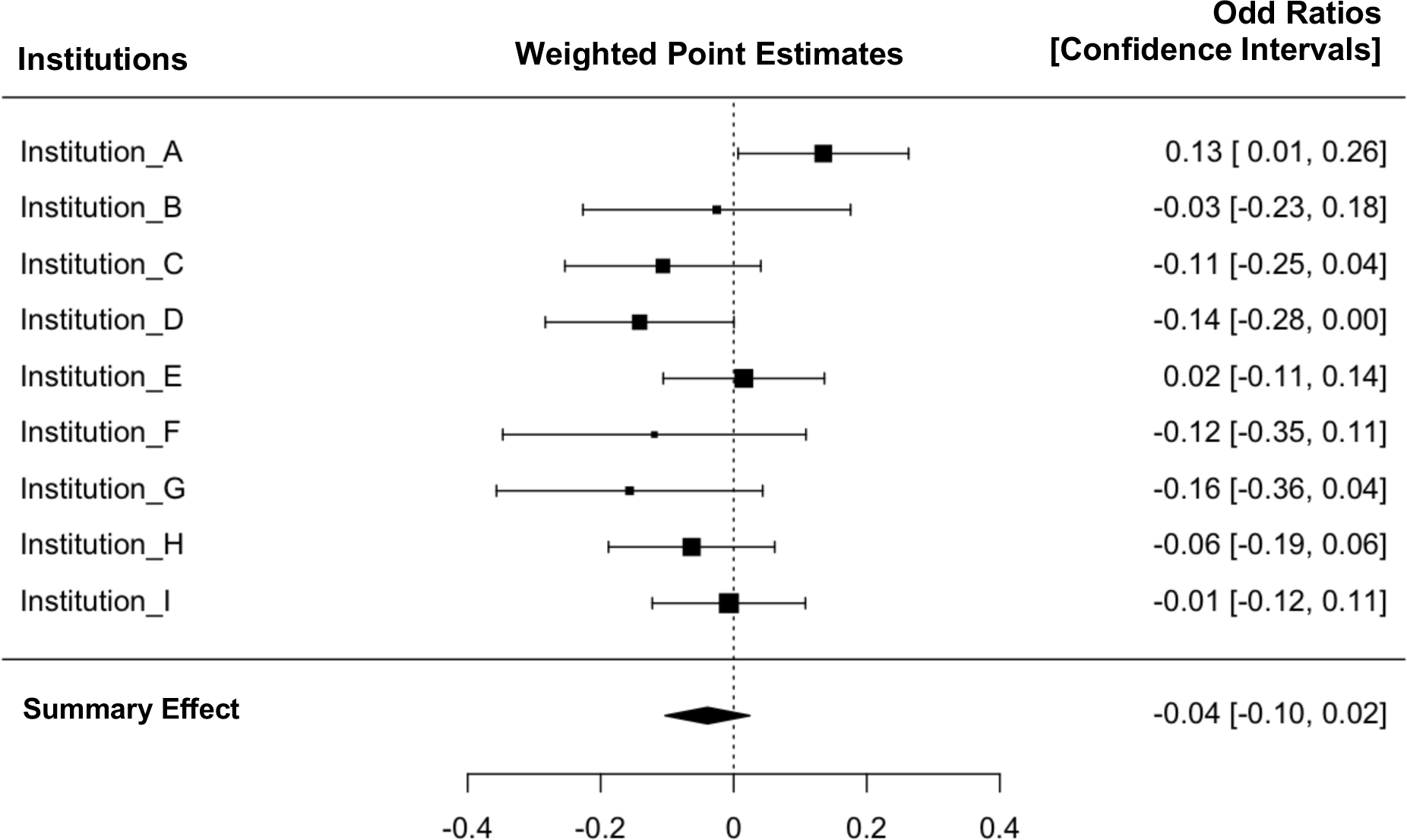
Graduate student productivity measured by total publications versus bivariate participation. Meta-analysis Forest plot displaying mean effect sizes (squares) and confidence intervals (brackets) for effect sizes of total publications versus bivariate professional development participation (control versus participants). Large squares denote greater impact on the summary effect based on sample size and effect size in each institutional sample. The vertical dotted line represents a null effect. The size and shape of the diamond at the bottom of the Forest plot represents the summary effect. Because the diamond overlaps the vertical line (null effect), this indicates that the effect of professional development participation on total publication is not significant.

**FIGURE 5.**
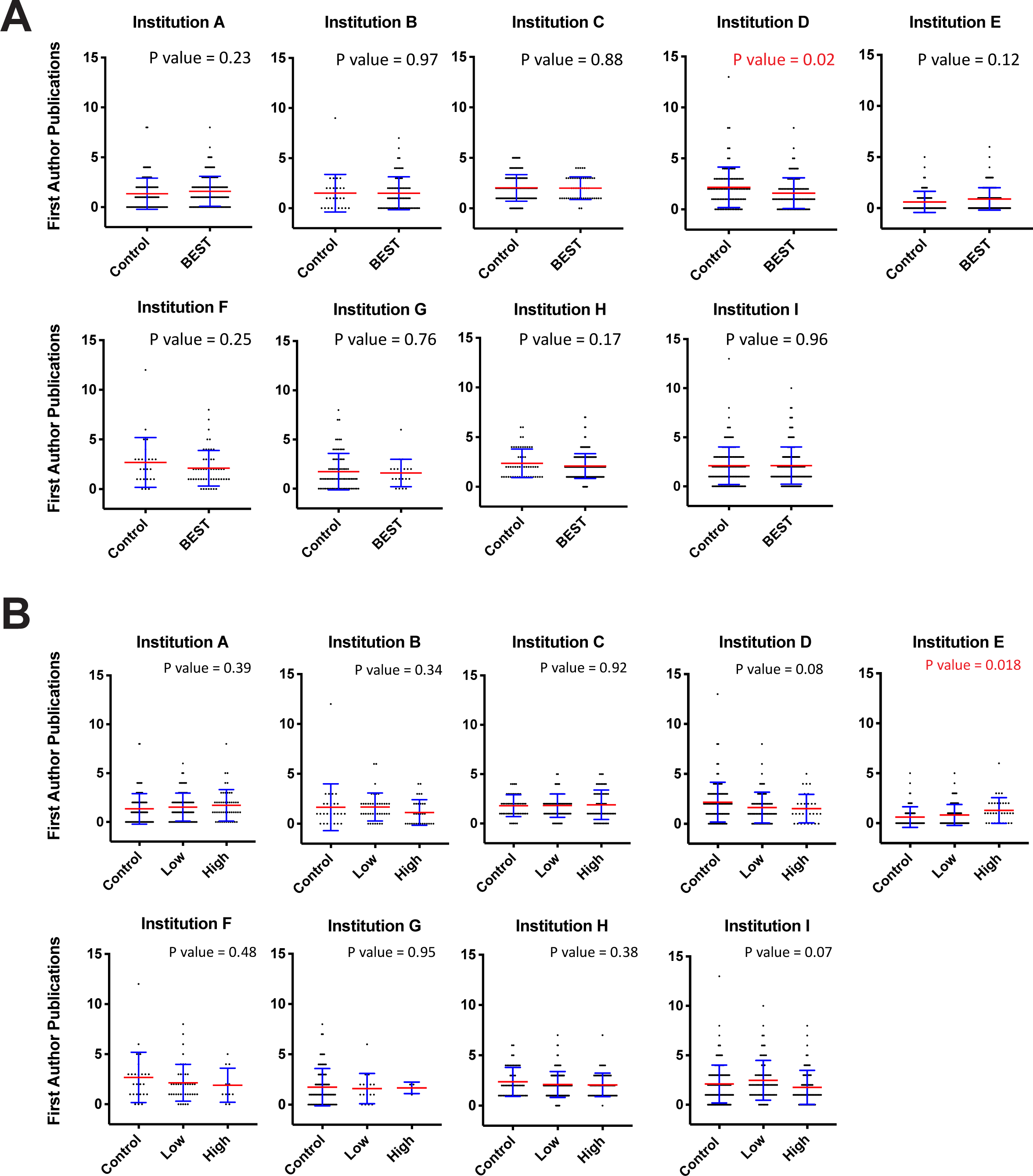
Professional Development Participation is Not Associated with a Decrease in First Author Publication. A, First author (including co-first author) publications versus binary professional development participation. Blue error bars represent standard deviation of the mean. Mean is denoted by a red line. Independent samples t-tests were used to compare of control vs. participant first author publications (significant values of p<.05 noted in red). B, First author (including co-first author) publications versus dosage of professional development participation. Blue error bars represent standard deviation of the mean. Mean is denoted by a red line. Analysis of variance was used to compare the impact of control, low, and high dose participation on first author publications (significant values of p<.05 noted in red).

**Figure 6.**
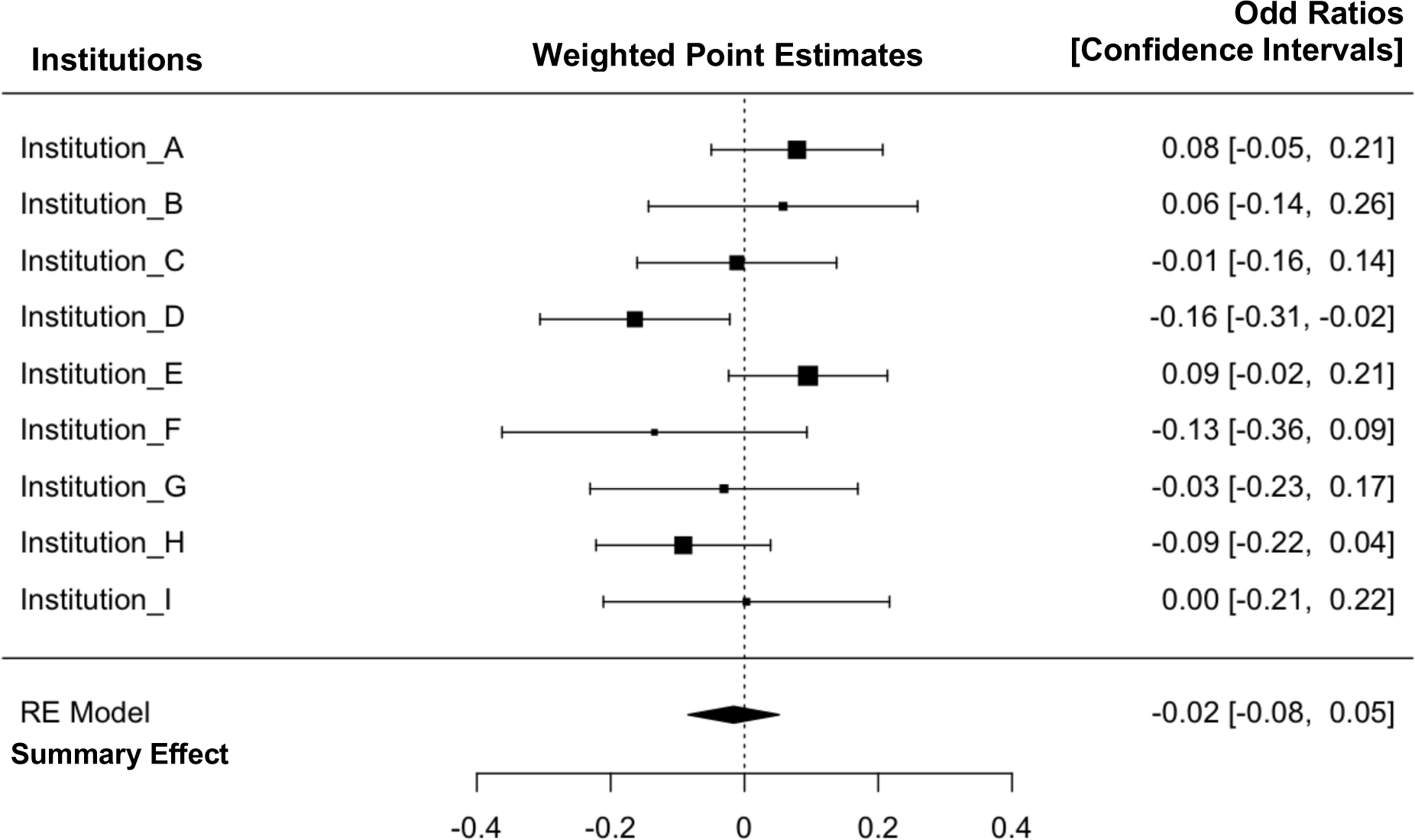
Graduate student productivity measured by first-author publications versus bivariate participation. Meta-analysis Forest plot displaying mean effect sizes (squares) and confidence intervals (brackets) for effect sizes of first-author publications versus bivariate professional development participation (control versus participants). Large squares denote greater impact on the summary effect based on sample size and effect size in each institutional sample. The vertical dotted line represents a null effect. The size and shape of the diamond at the bottom of the Forest plot represents the summary effect. Because the diamond overlaps the vertical line (null effect), this indicates that the effect of professional development participation on time to first author publications is not significant.

Power calculations suggest that with the average sample sizes and number of participating institutions’ data available for this analysis (N1=72, N2=115, alpha=0.05, k=9, *d*=0.20), and an observed moderate heterogeneity (I^2^ = 39.36% - 40.14% < 50% cutoff for moderate heterogeneity; τ^2^ =0.01, SEτ=0.01; Q=12.50 - 12.83, *p*=0.12-0.13) in a random effects model we have 87% power to detect a small effect size or larger (35, 36).

#### Weighted Publication Metric (PubMetric): An alternative comprehensive publication measure

Both first-author publications and total publications capture different aspects of productivity. By choosing to report one or the other, some information is lost. Instead of limiting the accuracy of reporting by removing one or the other, we proposed creating a novel publication metric that could capture trainees’ efforts on both types of contributions in a single metric. One concern that we anticipated was how to weigh these different contributions. For instance, first-author research papers may be valued over other types of contributions (*e.g*., middle-author research paper contributions or review papers). To address this issue, UNC developed a weighted publication metric (see Supplemental File 4, **S4**) that incorporates the four primary types of peer-reviewed publications into a single number. Impact factor was not included as a variable in the publication metric because impact factor as a measure of paper quality or journal prestige can be inherently biased by field. The UNC weighted publication metric was designed as a broader and more objective measure of the amount and quality of author contributions by trainees as reflected by authorship order.

To create the weighted publication metric, active training faculty at UNC were asked to rank the relative value of (A) first-author peer-reviewed <u>research</u> articles, (B) first-author peer-reviewed <u>review</u> articles, (C) middle-author peer-reviewed <u>research</u> articles, and (D) middle-author peer-reviewed <u>review</u> articles (n=150 responses from 350 total contacted; see **S4,** for details). First-author and co-first-author publications were considered synonymous. When averaging all faculty rankings and normalizing middle-author reviews to a weighting of 1, we generated the following equation for the weighted publication metric (PubMetric).

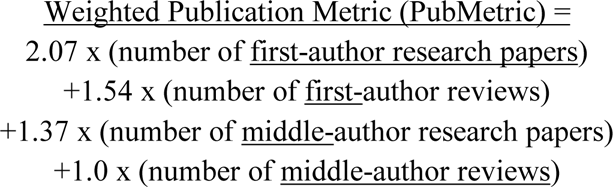

Four BEST institutions were able to provide weighted PubMetric data from PubMed scripts. Using this metric, similar patterns emerged as for total publications and first-author publications (**S4**).

### 4. Internships, Efficiency, & Productivity: Time to Degree, Total Publications, and First-author Publications

Internships are a form of career training that have unique characteristics and formats, but all require a relatively large time commitment that one could predict would impact time to degree or productivity (for definition of internship, see Supplemental File 3, **S3**). Institutions that supported internship opportunities provided outcome data for trainees who participated in their internship programs, which had differing lengths and designs and had some variant of a competitive selection process (**S3**).

We did not detect a difference in time to degree between graduate students who completed an internship and those who did not (**Figure 7a**). Similarly, we found no evidence of decrease in publication productivity or in the number of first-author publications in individuals that participated in an internship (**Figure 7b, 7c**). Internships were associated with favorable effect for some institutions’ first-author publications. Additional data on internship participation versus time to degree, total publications, first-author publications, and weighted publication metric showed no effect of participation (**S3**).

**FIGURE 7.**
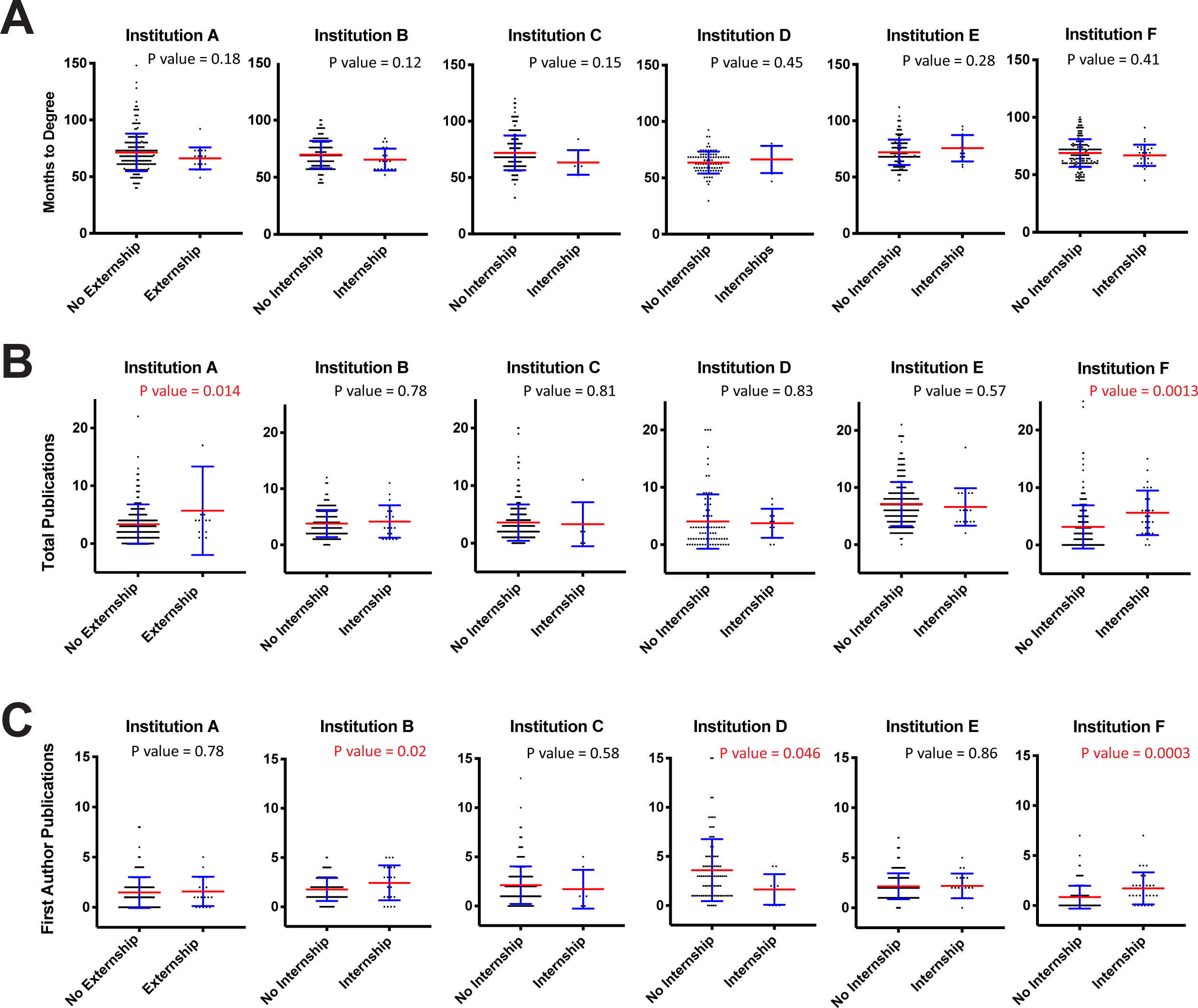
Internship or Externship Participation is Not Associated with an Increase in Time to Degree or a Decrease in Total or First Author Publications. A, Months to Degree versus internship or externship partici-pation. B, Total publications versus internship or externship participation. C, First author (including co-first author) publications versus internship or externship participation. Blue error bars represent standard deviation of the mean. Mean is denoted by a red line. Independent samples t-tests were used to compare of control vs. participant first author publications (significant values of p<.05 noted in red).

## Discussion

With concerns about productivity and length of doctoral education balanced with the need to provide adequate professional development, data from ten United States academic institutions were analyzed to determine if participation in career and professional development activities alters these outcomes. Here we discuss the impact of professional development on traditional metrics of academic success. Our study is unique in that it compiled doctoral degree durations at ten different universities, recorded individual participation in career and professional development activities in terms of dosage, and tracked individual engagement in real-time rather than relying on surveys sent to trainees after graduation.

The data show that even extensive participation did not result in a significant increase in time to degree or decrease in productivity of publications for doctoral graduate students in the life sciences. Overall, this is true for both low-dose and high-dose participants, although in our analyses, we found some significant changes in specific variables at some institutions.

Time to degree was chosen as a proxy for efficiency of completion because it was a measure at all institutions and facilitated comparisons. Publications were chosen as a proxy for productivity because they are an objective measure and because publications are widely viewed as an important currency of graduate performance in life science higher education (37, 38). The number of publications per graduate student in this study was in alignment with prior published work, where the average publication per graduate is 2.9 publications with a range of 0 to a maximum of 16 publications (39). Using our newly created weighted publication algorithm that considers all publications, the PubMetric, we found no difference in total number of publications between participants and controls at eight of the nine BEST institutions (**S4**).

Thus, across institutions nationwide, participating in career and professional development activities, including internships, did not negatively impact time to degree or manuscript publication. In fact, one institution even showed that participants with the highest dose (internships) had the most first-author publications. Although this observation could be partly explained by the fact that this program incorporated productivity into the selection process for internships, the same institutional requirements for first-author publications to graduate makes this explanation unlikely. Furthermore, other internship program institutions that recommended or required a first-author publication in order for a graduate intern to be selected also typically required one or more publications to graduate, reducing the likelihood that this explanation would fully account for the potentially beneficial effect.

## Limitations

Overall, a potential selection bias exists for the data because individual participants were not randomly selected but instead self-selected to participate (40). Some of this effect might be due to self-selection via the a la carte model, or program selection bias via an application-based cohort model. It is possible that these selected individuals were highly organized multi-taskers before participating and became better informed and motivated at BEST events.

One limitation to our cross-institutional comparison is that each BEST program independently defined what it meant to be a ‘participant’ in their program; similarly, definitions of control, low- or high-dose participation varied by program. Three institutions defined their dosage based on the number of hours of professional development; five institutions defined their dosage using the number of events attended; and two institutions grouped their participants by the number of credits or points assigned for attendance.

Just as the program offerings of each institution were unique, so too were the trainee populations that were eligible for programming (**S1**). Some BEST institutions required trainees to apply to the program and participate in activities as a cohort while other BEST institutions used an a la carte model so that trainees could choose from among professional development offerings.

Others used a combination of cohort and a la carte, and some gradually opened program activities to more participants due to demand. For this reason, a classic “control” population (i.e. zero participation in professional development activities) is difficult to define when evaluating the impact of BEST programs. In addition, even the “control” population may have participated in other professional development events sponsored by other campus offices or student groups, scientific societies, companies, or other external organizations.

## Culture Change

Notably, the US government clarified that researchers in doctoral and postdoctoral training who are supported by any federal funds are expected to not only conduct their research, but are also allowed to devote time to career and professional development (41). These guidelines helped faculty to better accept the notion of doctoral students’ participation in activities outside of dissertation research.

Studies published by BEST institutions have further reinforced this change in faculty attitudes (25). These studies showed that faculty’s initial hesitation is evolving to an understanding that next-generation scientists will not only need to be excellent researchers, but also need to be equipped with professional skills that are more effectively learned outside the laboratory. This viewpoint is supported by a snapshot of current faculty perceptions, which was obtained using subgroup surveys launched by institutions receiving NIH BEST funding. Responses showed that faculty believe that BEST career development programming is beneficial to trainees in a number of different ways: no delayed time to degree, enhanced happiness, positive effects in the lab, and more confidence in directing trainees’ own career development (25).

Our current study, based on quantitative data, supports that participation in BEST programming did not adversely affect time to degree or numbers of manuscripts published, and in select cases, even correlated with a shorter or more productive outcome. We predict that as more evidence-based support for professional development comes to light, more faculty members will feel confident in encouraging their students to participate in such programming. Although further studies are needed to extend these conclusions across disciplines, the authors hope that these data will assuage concerns of faculty and trainees alike. Historically, biomedical sciences faculty had expressed initial hesitation toward time spent in first year laboratory rotations when first instituted, yet doctoral time to degree tracking at Cornell University revealed no statistically significant lengthening across comparison groups before and after rotations were mandated in 2003 for three graduate fields (**S1**) *–* and rotations are now a widely accepted best practice within the biomedical sciences. We hope that similarly, our data will provide encouragement for professional development during training to become equally commonplace as an accepted foundation of PhD training.

Over the past decade, academic institutions have increasingly recognized the breadth of careers pursued by doctoral students and the need for interventions and resources to support their future success (42). Many institutions have rapidly incorporated career and professional development training within doctoral programming (43, 44). Transformational training programs are well positioned to flourish because of this new training environment, heightened faculty awareness, institutional commitment, and support from funding agencies. Such examples are found at The Burroughs Wellcome Fund with its Career Guidance for Trainees Grant, the NIH National Institute of General Medical Sciences with its Innovative Programs to Enhance Research Training and Career Development Supplement, and the National Science Foundation with its Research Training Programs (11–13, 42, 45).

We hope that readers will share the results of our current study with their colleagues, and incorporate experiential learning activities into PhD training programs (3, 46, 48, 50). Although this study focused on doctoral students from biomedical fields, we anticipate that the major conclusions of this study are likely applicable to graduate students and postdoctoral researchers in other STEM fields, as well as to other fields including those in the humanities, arts and social sciences.

## Acknowledgements

The authors would like thank Christopher Pena and Michael Friedlander for support of their institutional BEST projects; Jacek Buczny for advising on data analysis; and Patricia Labowski for her continual support of the BEST institutions, its investigators, and her leadership of the larger national initiative. We thank the Graduate Career Consortium Graduate Education Research Committee for helpful comments on the manuscript. Funding sources included the Common Fund NIH Director’s Biomedical Research Workforce Innovation Broadening Experiences in Scientific Training (BEST) Award (DP7OD020317). The following institutional NIH BEST awards (alphabetical by institution) included: DP7OD020322 (Boston University); DP7OD020316 (Chicago); DP7OD018425 (Cornell University); DP7OD018428 (VA Tech); DP7OD020314 (Rutgers); (Rochester); DP7OD020317 (Vanderbilt); DP7OD018424 (University of California, Irvine); DP7OD020317 (University of North Carolina, Chapel Hill); DP7 OD018427 (Wayne State). National Institutes of Health (NIH) General Medical Sciences - Science of Science Policy Approach to Analyzing and Innovating the Biomedical Research Enterprise (SCISIPBIO) Award (GM-19-011) - 1R01GM140282-01 (UNC).

## Author Contributions

*Conceptualization* - ABS, AM, AMB, AVW, BMS, BB, CAS, CSC, DA, DF, HS, ID, JA, JDH, KAP, LH, MF, PDB, PJB, RC, RLL, SSV, TB

*Data curation* - ABS, AMB, AVW, CAS, CSC, JA, HS, ID, KAP, PDB, RLL, SSV, TB

*Formal analysis* - PDB, PJB, RLL

*Funding acquisition* - AM, AVW, BB, BMS, CSC, DF, HS, KAP, KLG, LH, MF, PDB, PJB, RC, RLL, SSV

*Investigation* - PDB, PJB, RLL

*Methodology* - PDB, PJB, RLL

*Project Administration* – ABS, AM, AMB, AVW, CAS, CSC, HS, JA, KP, PDB, RLL, SSV, TB

*Software* – DA, JDH

*Supervision* - PDB, PJB, RLL, TB

*Validation* - ABS, AMB, AVW, CAS, CSC, DA, HS, ID, JA, JDH, KAP, PDB, RLL, SSV, TB

*Visualization* - PDB, PJB, RLL, SSV

*Writing – Original Draft Preparation* - ABS, ID, PDB, RLL, PJB, SSV, TB

*Writing – Review & Editing* - ABS, AMB, AM, AVW, BB, BMS, CAS CEH, CSC, DA, DF, HS, ID, JA, JDH, KAP, KLG, LH, MF, PDB, PJB, RC, RLL, SSV, TB

## SUPPLEMENTAL INFORMATION

### Supplemental File 1. BEST Institutional and Program Details

**SI Table 1a.**
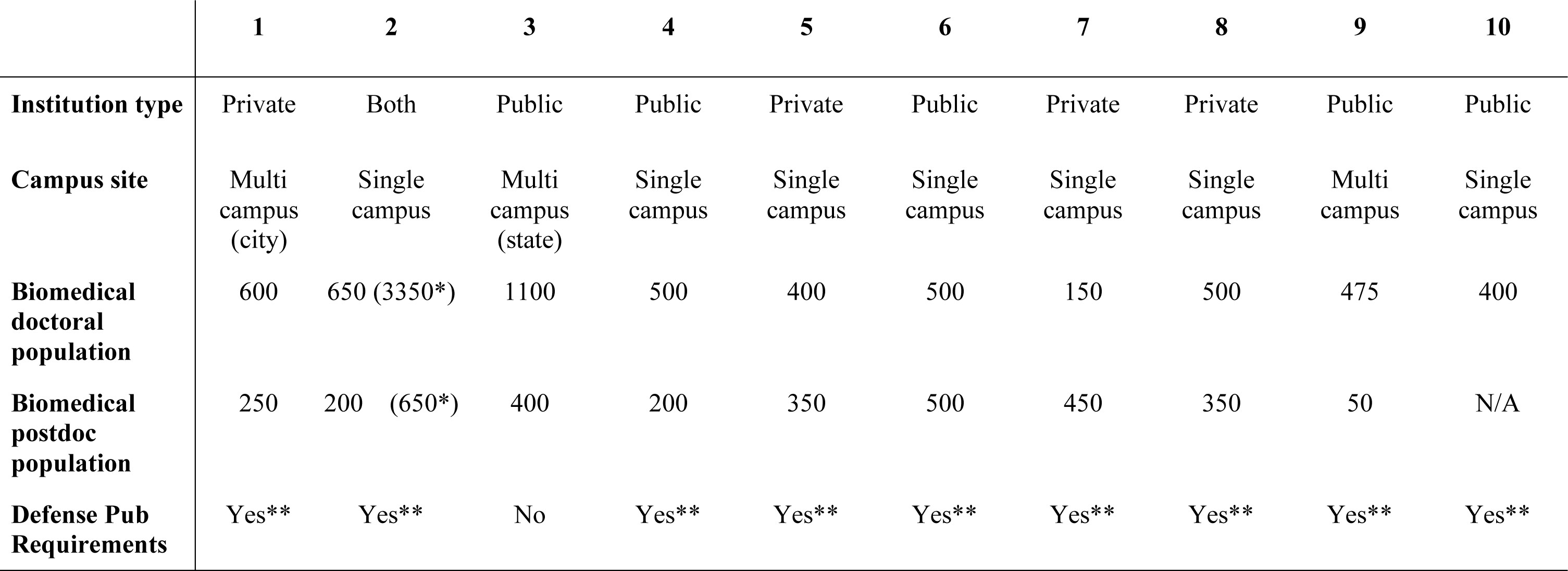
Institutional Profiles **SI Table 1a Legend.** * Trainees across all discipline. Campuses located across multiple sites, city- or state-wide, are indicated. ** Program dependent. *** Grad school requirements. *Note*: all institutions (appear in alphabetical order): 1) Boston University, 2) Cornell University, 3) Rutgers University, 4) University of California, Irvine, 5) University of Chicago, 6) University of North Carolina at Chapel Hill, 7) University of Rochester, 8) Vanderbilt University, 9) Virginia Tech, and 10) Wayne State University.

**SI Table 1b.**
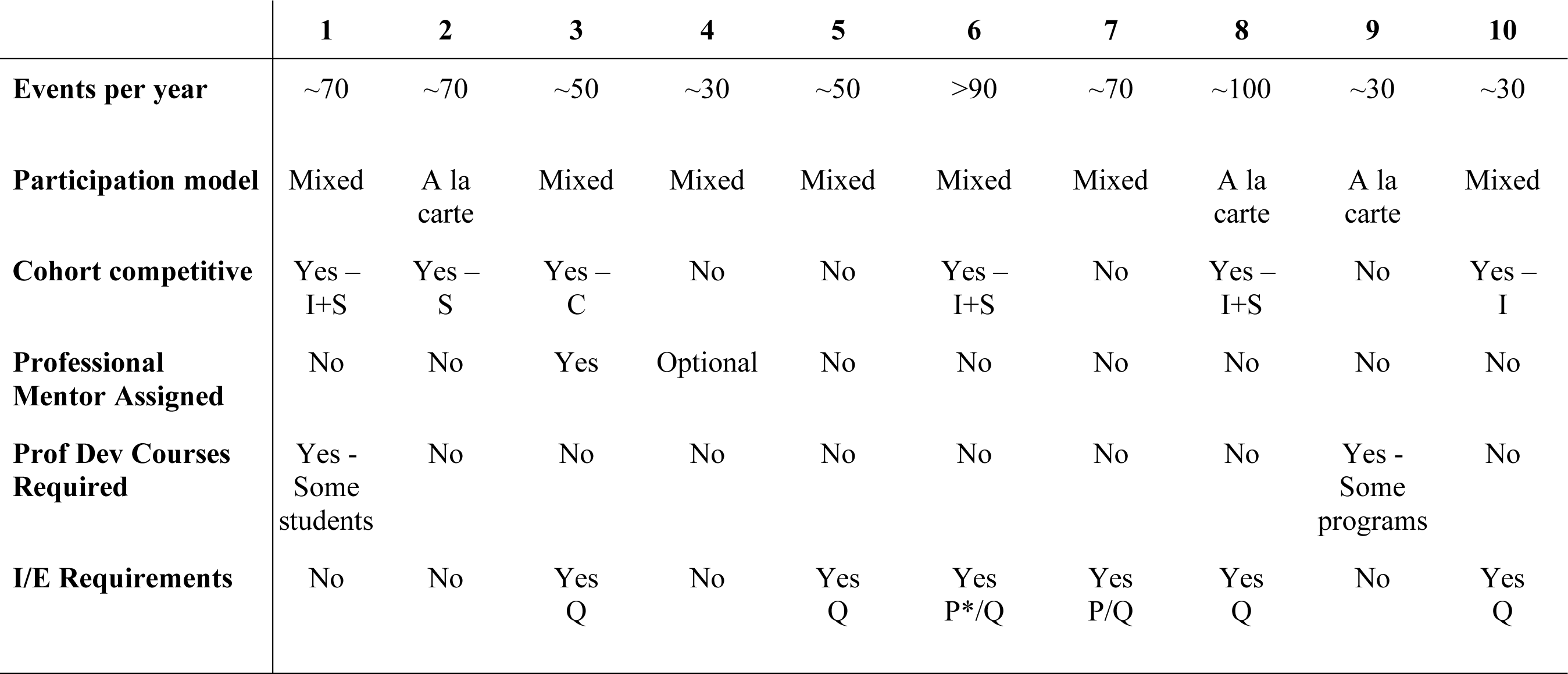
BEST Program Activities and Participating Departments **SI Table 1b Legend.** I/E= internship or externship. Q = Qualifying Exams required for internship/externship, P= Publications required for internship/externship, P*= first-author publication recommended. “Participation model” refers to whether the BEST programs were open to all trainees (a la carte) or whether BEST programs were open to only a selected group of trainees (cohort). I+S= competitive for internships and site visits, I, S= competitive for internships or site visits only; C competitive for cohort; N/A = not applicable. Where offered, internships were always optional (opt-in) activities, as were externships, shadowing, and site visits, whereas professional development course were required by some. *Note*: all institutions (appear in alphabetical order): 1) Boston University, 2) Cornell University, 3) Rutgers University, 4) University of California, Irvine, 5) University of Chicago, 6) University of North Carolina at Chapel Hill, 7) University of Rochester, 8) Vanderbilt University, 9) Virginia Tech, and 10) Wayne State University.

**SI Table 1c.**
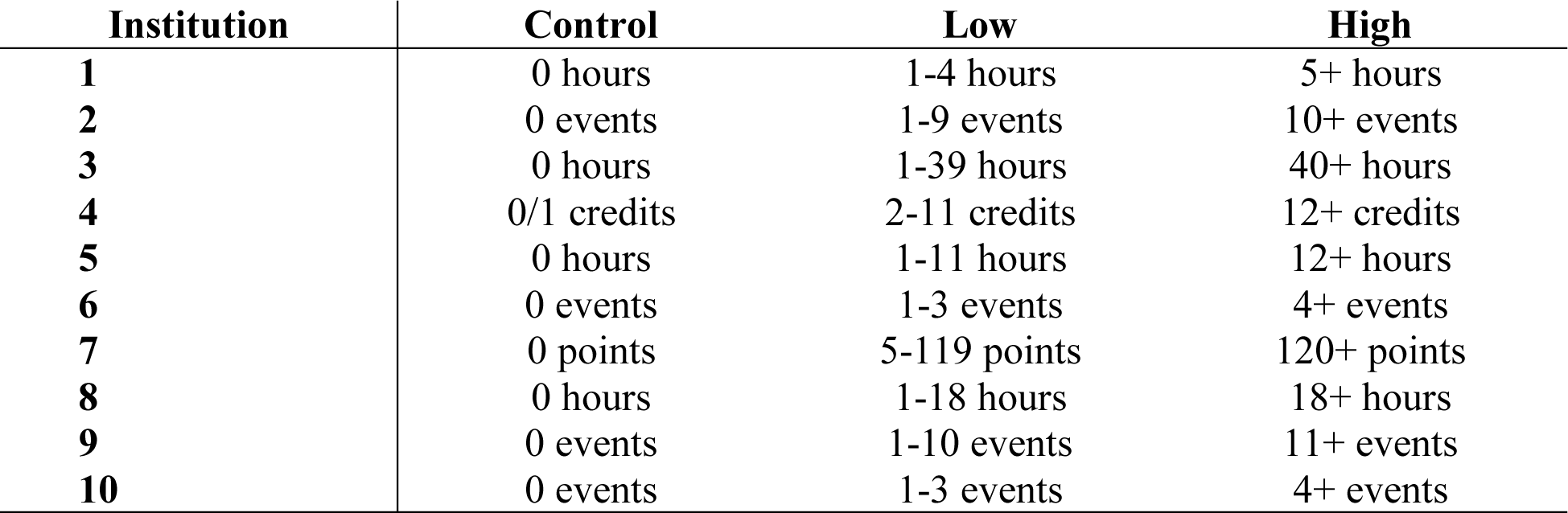
**Definition of dosage SI Table 1c Legend.** Participation was recorded at each institution as hours, events, or points. All bivariate analyses contrast control with any dosage (low plus high dosages combined). All dose-response analyses use the grouped definitions for control, low, and high as noted.

**SI Figure 1.**
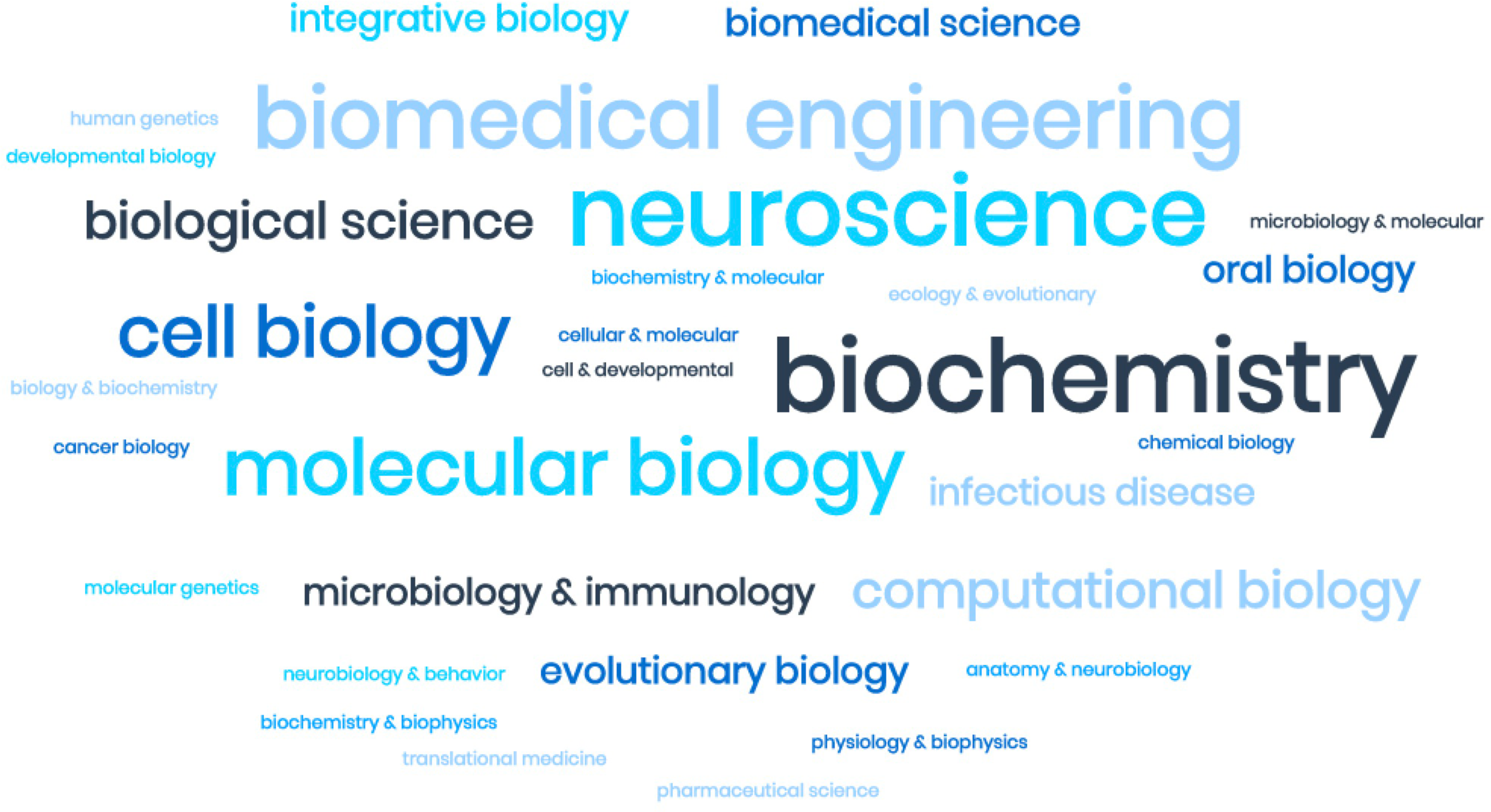
Visualization of common departments included in sample: Word cloud generator of participating departments

**SI Table 1d.** Graduate programs/departments represented in each institution’s dataset (listed alphabetically by institution)

| Institution | Departments, Programs, & Fields |  |
| --- | --- | --- |
| **Cornell University:**<br>*Graduate Fields (n=14)* | ▪ Biochemistry, Molecular & Cell Biology<br>▪ Biological & Environmental Engineering<br>▪ Biomedical Engineering<br>▪ Biophysics<br>▪ Biomedical & Biological Sciences<br>▪ Computational Biology<br>▪ Ecology and Evolutionary Biology | ▪ Genetics and Development<br>▪ Immunology and Infectious Disease<br>▪ Microbiology<br>▪ Molecular and Integrative Physiology<br>▪ Neurobiology and Behavior<br>▪ Nutrition |
| **Boston University:**<br>*Programs (n=19)* | ▪ Anatomy & Neurobiology<br>▪ Behavioral Neuroscience<br>▪ Biochemistry<br>▪ Bioinformatics<br>▪ Biology<br>▪ Biomedical Engineering<br>▪ Biophysics<br>▪ Brain, Behavior, and Cognition<br>▪ Chemistry<br>▪ Genetics & Genomics | ▪ Graduate Program for Neuroscience<br>▪ Microbiology<br>▪ Molecular, Cell Biology & Biochemistry<br>▪ Molecular & Translational Medicine<br>▪ Nutrition & Metabolism<br>▪ Oral Biology<br>▪ Pathology & Laboratory Medicine<br>▪ Pharmacology & Experimental Therapeutics<br>▪ Physiology<br>▪ Program in Biomedical Sciences |
| **University of North Carolina, Chapel Hill:**<br>*Departments, Programs, & Curricula (n=16)* | ▪ Bioinformatics and Computational Biology<br>▪ Biochemistry and Biophysics<br>▪ Biology<br>▪ Biomedical Engineering<br>▪ Cell Biology and Physiology<br>▪ Chemistry<br>▪ Genetics and Molecular Biology<br>▪ Microbiology and Immunology | ▪ Neuroscience<br>▪ Nutrition<br>▪ Oral Biology<br>▪ Pathobiology and Translational Medicine<br>▪ Pharmacology<br>▪ Chemical Biology & Medicinal Chemistry<br>▪ Molecular Pharmaceutics<br>▪ Toxicology |
| **University of California, Irvine:**<br>*Departments (n=16)* | ▪ Anatomy and Neurobiology<br>▪ Biological Chemistry<br>▪ Biomedical Engineering | ▪ Experimental Pathology<br>▪ Psychiatry & Human Behavior<br>▪ Microbiology and Molecular Genetics |
|  | ▪ Chemical Engineering<br>▪ Chemistry<br>▪ Developmental & Cell Biology<br>▪ Ecology & Evolutionary Biology<br>▪ Epidemiology | ▪ Molecular Biology & Biochemistry<br>▪ Neurobiology & Behavior<br>▪ Pharmacological Sci<br>▪ Physiology and Biophysics<br>▪ Public Health |
| **Virginia Polytechnic Institute and State University:**<br>*Departments and Programs (n=14)* | ▪ Animal and Poultry Sciences<br>▪ Biochemistry<br>▪ Biological Sciences<br>▪ Biomedical Sciences & Pathobiology<br>▪ Human Development<br>▪ Human Nutrition, Foods, and Exercise<br>▪ Psychology<br>▪ School of Biomedical Engineering and Sciences | ▪ Translational Biology, Medicine, and Health:<br>▪ Cancer<br>▪ Development, Aging, and Repair<br>▪ Health Implementation Science<br>▪ Immunity and Infectious Disease<br>▪ Metabolic and Cardiovascular Science<br>▪ Neuroscience |
| **Rutgers University:**<br>*Departments (n=25)* | ▪ Cell Biology, Neuroscience and Physiology<br>▪ Biomedical Engineering<br>▪ Infection, Immunity and Immunology<br>▪ Molecular Biology, Genetics and Cancer<br>▪ Oral Biology<br>▪ Biochemistry<br>▪ Cell & Developmental Biology<br>▪ Exposure Science<br>▪ Microbiology & Molecular Genetics<br>▪ Neuroscience<br>▪ Pharmacology, Cellular & Molecular<br>▪ Physiology & Integrative Biology | ▪ Toxicology<br>▪ Chemical & Biochemical Engineering<br>▪ Chemistry & Chemical Biology<br>▪ Computational Biology & Molecular Biophysics<br>▪ Endocrinology Sciences<br>▪ Medicinal Chemistry<br>▪ Pharmaceutical Sciences<br>▪ Microbial biology<br>▪ Nutritional Sciences<br>▪ Behavioral & Neural Sciences<br>▪ Biology<br>▪ Environmental Sciences<br>▪ Computational & Integrative Biology |
| **University of Rochester:**<br>*Departments (n=15)* | ▪ Translational<br>▪ Biomedical Sciences<br>▪ Biomedical Engineering | ▪ Cardiovascular Research Institute<br>▪ Ophthalmology<br>▪ Environmental Medicine (Toxicology) |
|  | ▪ Microbiology and Immunology<br>▪ Neurosciences and Neurobiology and Anatomy<br>▪ Biochemistry & Biophysics<br>▪ Center for Musculoskeletal Research<br>▪ Brain and Cognitive Sciences | ▪ Biomedical Genetics<br>▪ Pediatrics Infectious Diseases<br>▪ Pharmacology and Physiology<br>▪ Biostatistics and Computational Biology<br>▪ Pathology |
| **Vanderbilt:**<br>*Programs & Departments*<br>*(n=11)* | ▪ Biochemistry<br>▪ Biological Sciences<br>▪ Cancer Biology<br>▪ Cell & Developmental Biology<br>▪ Cellular & Molecular Pathology<br>▪ Chemical & Physical Biology | ▪ Human Genetics<br>▪ Microbiology & Immunology<br>▪ Molecular Physiology & Biophysics<br>▪ Neuroscience<br>▪ Pharmacology |
| **Wayne State:**<br>*Programs & Departments*<br>*(n=12)* | ▪ Biochemistry and Molecular Biology<br>▪ Biological Sciences<br>▪ Biomedical Engineering<br>▪ Chemistry<br>▪ Immunology and Microbiology<br>▪ Molecular Biology and Genetics | ▪ Nutrition and Food Science<br>▪ Pathology<br>▪ Pharmaceutical Sciences<br>▪ Pharmacology<br>▪ Physiology<br>▪ Translational Neuroscience |
| **University of Chicago:**<br>*Graduate programs*<br>*(n=17)* | ▪ Biochemistry & Molecular Biology<br>▪ Cancer Biology<br>▪ Cell and Molecular Biology<br>▪ Computational Neuroscience<br>▪ Development, Regeneration & Stem Cell Biology<br>▪ Ecology and Evolution<br>▪ Evolutionary Biology<br>▪ Genetics, Genomics and Systems Biology | ▪ Public Health Sciences<br>▪ Human Genetics<br>▪ Immunology<br>▪ Integrative Biology<br>▪ Medical Physics<br>▪ Microbiology<br>▪ Molecular Metabolism and Nutrition<br>▪ Molecular Pathogenesis and Molecular Medicine<br>▪ Neurobiology |

**SI Table 1e.** **NIH BEST Programming & Awardee Program Descriptions**

| **Organization/**<br>**Institution** | **Description** |
| --- | --- |
| ***NIH BEST Program:*** | **Development of Programming Definitions:** Despite common data collection forms, collection procedures, and reporting methods, some inconsistencies between programs’ definitions remained, thus resulting in the convening of the BEST Consortium Data Summit, with the shared goal of establishing definitions of terms that were used in our common NIH BEST/Windrose reporting requirements. For example, there was initially confusion over what constituted a ‘workshop’ versus a ‘seminar’; consensus established at the Data Summit clarified a workshop as having a customized product or take-home material (*e.g.*, personal answers to interview questions, results of a job simulation exercise, goal list, draft individual development plans; IDPs), whereas a ‘seminar’ was defined as passive listening to content delivery. Shared definitions of key terms enabled common data comparison/collection by the NIH (Lenzi et al, 2020) and have enabled collaboration between BEST institutions to compare and publish significant program outcomes in reports like this one. |
| ***Boston University BEST Program:*** | BU’s BEST is based on a classic feedback loop. Job market analysis informs program development which in turn equips trainees with the skills required to successfully join the workforce. Using the Labor•Insight™ software tool developed by Burning Glass Technologies, jobs, job trends, job locations, and so-called “hard” and “soft” skills required for various career pathways are revealed. With this information in hand, BU's BEST offers activities to equip trainees with skills needed in six broad biomedical career tracks. The programming is designed to enable the trainees to reflect on their career interests, to explore various career paths and to enhance their skills to prepare for a productive career. Trainees are encouraged to work on an IDP to gain insight into future career possibilities while participating in coursework, workshops and panel discussions with local professionals to gain additional knowledge about options. Once trainees’ interests are refined to a particular career track, they can participate in offerings to hone the skills identified by Labor•Insight™. Examples of workshops include those dedicated to grant writing, data analysis, entrepreneurship, creating a successful resume, LinkedIn profile or cover letter. Site visits and internships are also offered for more experiential learning. Finally, one-on-one career coaching is available and our trainees are encouraged to utilize our alumni mentor network for informational interviews. Taken together, these tools help trainees prepare to pursue their chosen path. |
| ***Cornell University***<br>***Careers Beyond***<br>***Academia/BEST:*** | Cornell University's Careers Beyond Academia/BEST provides flexible, experiential, empowering personalized opportunities for doctoral students and postdocs in all disciplines to make informed choices about their careers. Exposure to career options for PhDs comes via seminars in collaboration with department series organizers, workshops, signature "Careers in..." panels, symposia, employer site visits and courses to provide hands-on experience, all with an underlying mentoring component. Partnerships with several trainee-run associations formed and supported include the Cornell Graduate Consulting Club (CGCC), Advancing Science and Policy (ASAP), the Technology & Entrepreneurship Club (TEC), Engineering Graduate Student Association (EGSA), Chemical Biology Interface (CBI training grant) and additional student-run organizations to address programming gaps based on iterative feedback. All together we provide group and individual coaching/advising sessions, training, interactions with practitioners, case competitions, practice describing their expertise in the language of their future employer (or funder) orally and online, and practical advice on researching and obtaining a job using the skills learned. Students and postdocs are awarded funds by application to attend conferences, join professional societies beyond their academic discipline, and create their own activities which often involve alumni. Embedded in the Graduate School, we partner with the Office of Postdoctoral Studies, Entrepreneurship@Cornell, the Society for Humanities, Career Services, CU-CIRTL, the Center for Teaching Innovation and other on-campus groups to cover additional focus areas that we co-develop and co-advertise. Careers Beyond Academia/BEST enhances training opportunities for graduate students and postdoctoral scholars in all fields through an individualized, flexible program that empowers trainees to acquire the knowledge and hone skills to become more credible for an ideal career outcome. KEYS to SUCCESSES. Flexibility: a program that is ready when students and postdocs are, offering opportunities at the dose they are ready to receive, increases both faculty buy-in and trainee empowerment. Personalization: as no two careers are identical, neither will their training needs be so; we also encourage and support student/postdoc-initiated ideas. Experiential opportunities: to be able to say 'I've done that' and write it on their resume. Gaining professorial buy-in: it is dependent on how we have marketed the program the skills learned will foster success in any field, including academia we are not pushing students to careers beyond academia but rather are enabling an informed choice for future success. We offer a resource to alleviate faculty pain points (if they have no experience in industry, or feel they can't connect trainees with mentors or opportunities in science policy or intellectual property law, e.g.) and to showcase their successful alumni. |
| ***Rutgers iJOBS***<br>***Program:*** | iJOBS (interdisciplinary Job Opportunities for Biomedical Scientists) consists of four phases and is designed to be very flexible. Phase 1 is open to all trainees and consists of weekly a la carte events to inform trainees of the various career options. Activities include hands-on job simulations, half day site visits to companies, |
|  | skill primers, career panels, workshops, career fairs, and a 4-day intensive workshop SciPhD: Leadership and Business Skills for Scientists. Trainees who are interested in doing a deeper dive into a particular career track can apply to the Phase 2 cohort once they have completed at least 12 hours of Phase 1 events. About 20 trainees per year are admitted to Phase 2 and are allowed to take part in intensive training and coursework for their chosen career track. Specifically, industrial partners host unpaid shadowing/externships to allow for real world exposure (total of 72 hours over the course of a semester for observational purposes only). Trainees also register for a course to increase their knowledge base (e.g. business or law school class), and complete Individual Development Plans (IDPs) with guidance from a professional mentor. During Phase 3, trainees prepare for job search and placement via professional coaching on their resume, LinkedIn profile, and interview skills. Upon successful career placement, iJOBS program alumni are encouraged to share their wisdom with new iJOBS trainees in Phase 4 as they serve as mentors, event hosts, and shadow partners. |
| ***University of Chicago myCHOICE Program:*** | The myCHOICE program benefits from a broad-based steering committee that includes representatives from the UChicago Biological Sciences Division (BSD) postdoctoral association, graduate student dean's council, and diversity committee, and from the UChicago biotech association, Center for teaching, Graduate Student Affairs office and Provost's office. The myCHOICE program is open to all graduate students and postdoctoral scientists pursuing biological sciences training at UChicago. Programming emphasizes both career exposure and professional development. Career exposure covers ten general areas based on the myIDP (Individual Development Plan from Science Careers) categories. Exposure areas include such broad topics as Industry, Tech Commercialization, Entrepreneurship, Medicine/Healthcare, Business of Science, Law, Communication, Teaching, and Administration, and also provides exposure to Academic Research. myCHOICE is characterized by a three-phase training plan based around Exposure, Education and Experience (E1-E3) to the ten career areas of the above topics. The E1: Exposure seminar series "What can I do with my PhD" is open to all participants including those from other institutions. E2 and E3 level programming requires a myCHOICE application, including completion of the myIDP. UChicagoOverview: Among the unique features of the myCHOICE program also are the inclusion of guided mentorship from alumni and friends of the institution. and "outstanding leveraging of internal and external resources and the experimental nature of the plan with hypothesis testing" (quoted directly from the Resume and Summary of Discussion from peer review). Internal resources include the Booth School of Business, the Harris School of Public Policy, UChicagoTech, and the Chicago Innovation Exchange (the location of some myCHOICE events). External resources include MATTER (a Chicago based healthcare innovation center), the Alan Alda Center for Communicating Science and several local industry partners. myCHOICE benefits from a broad-based steering committee that includes representatives from the UChicago Biological Sciences Division |
|  | (BSD) postdoctoral association, graduate student dean's council, and diversity committee, and from the UChicago biotech association, Center for teaching, Graduate Student Affairs office and Provost's office. The myCHOICE innovative evaluation plan is designed to test two hypotheses. More extensive participation in myCHOICE predicts greater PhD and postdoctoral scientist career choice empowerment and satisfaction with chosen career. More extensive participation in myCHOICE predicts improved concordance between myIDP Career Fit assessment at training exit and actual career selection. A survey of UChicago BSD graduate students after just one quarter (3 months) of myCHOICE programming has indicated that 90% of students are aware of the program and 70% have already attended at least one event. |
| ***UC Irvine GPS-BIOMED Program:*** | UCI's Graduate Professional Success in the Biomedical Sciences (GPS-BIOMED) aims to better prepare graduate students and postdoctoral scholars for a variety of careers within the biomedical research workforce, and empower trainees to become not only skilled scientists, but also polished professionals. GPS-BIOMED is a campus-wide program spanning four schools and 20+ departments and PhD programs. Our program has a 4 pillar model: **Explore:** Increase awareness and interest in diverse science-related careers via Career nights & Life Beyond PhD seminar series. **Train:** Improve professional skills required for success in diverse fields. Science Communication Skills and Extension courses help trainees broaden the scope of career training. **Experience:** Provide hands-on experience through industry site visits, internships and externships. We offer on-campus internships as well as off-campus opportunities within local companies including, Allergan, Edwards Lifesciences, Medtronic and Evoke Neurosciences. **Transition:** Build networks that allow students and postdocs to prepare for and transition to science-related careers. Mentorship and networking are key elements for successful career transitions. With help from external advisors we host networking mixers where visitors from different job sectors discuss opportunities and daily job functions. We also have a one-on-one mentoring program. Our program has also added value to research efforts at UCI e.g. improved scores on training grants and fellowships and new partnerships with industry. The program has helped with recruitment of talent pool. 30% of the incoming students mentioned that the presence of GPS-BIOMED program helped them make a positive decision to join UCI. Additionally, based on the alumni survey, a large percentage of them owe their career preparedness and success to GPS-BIOMED program, increasing alumni engagement. |
| **University of North Carolina at Chapel Hill TIBBS Program:** | The Training Initiatives in Biological and Biomedical Sciences (TIBBS) programming supplements our trainees scientific training with the non-bench skills needed to be successful in a wide range of careers through: regular professional development programming, student-led career cohorts, workshop series, site visits, and an immersive internship program. Professional development workshops (e.g., fellowship writing, career planning) supplements the career exposure available through the career cohorts an immersive |
|  | experiences. TIBBS plans events and workshops targeted to doctoral and postdoctoral scientists at specific career stages, including TIBBS sponsors the Annual Career Blitz, which brings two-dozen scientists from a wide variety of research and research-related careers in and out of academia, to campus for an afternoon of instruction and networking (typically attendees by nearly 200 scientists annually). Career cohorts focus on a wide variety of career pathways, including: business and consulting; science policy and outreach; writing and communication; teaching intensive careers; and academic and research-intensive careers. TIBBS provides structure and support for scientist-led career cohorts that meet monthly to network with invited professionals, share career resources, and report back on informational interviews. Doctoral and postdoctoral scientists gain leadership experience through their groups and groups frequently collaborate to bring in external scientists whose job duties span interest areas. Workshop series take place two to three times per year, alternating topics of interest, mirroring each of the cohort's popular interest areas (e.g., policy series communication series, teaching series, and research-intensive careers series), as well Nationally known consultants and speakers anchor the workshop series that are supplemented by expert local knowledge. as including topics spanning career areas (e.g., leadership series, and industry skills series). UNC's proximity to the Research Triangle Park situates us to take advantage of multiple immersive learning opportunities, including site visits, and internships. Doctoral and postdoctoral scientists go on monthly field trips to local companies, organizations, and non-profits representing nearly all available career options, co-organized with local institutional partners (e.g., ELITE Consortium). TIBBS represents UNC's long-standing and continuing commitment to professional and career development for our 1000 biological and biomedical graduate student and postdoctoral scientists. |
| ***University of Rochester URBEST Program:*** | UR's Broadening Experiences in Scientific Training (URBEST) program funds instruction in leadership and professionalism, creates new opportunities for experiential learning through internships and shadowing, and provides training pathways in (1) industry, manufacturing and entrepreneurship; (2) regulatory affairs, compliance and review; and (3) science and technology policy. First URBEST program activities include an Individual Development Plan (IDP) Workshop, a Leadership Advantage Program, and a URBEST Retreat and Career Workshop. Internship opportunities are available to select URBEST participants who show research productivity and career development initiative. Novel components of the URBEST program include: Incorporating Self-Determination Theory (SDT) into IDPs and program evaluations to ensure continuous improvement and effectiveness of URBEST activities. Faculty, alumni and peer mentoring networks have been established using a URBEST LinkedIn Group and SDT-based Mentoring Workshops to foster improved mentoring practices and better support scientist autonomy and diverse career outcomes. Other activities include a flipped-classroom Career Stories Q&A Seminar series, and new courses tailored for URBEST |
|  | participants, such as an Intellectual Property and Commercializing Technology Seminar Series and a Leadership and Management for Scientists Course. |
| **Vanderbilt ASPIRE Program:** | The ASPIRE Program is designed to empower Vanderbilt’s biomedical sciences PhD students and postdoctoral scholars to make well-informed career decisions with confidence. ASPIRE provides PhD students and postdocs (collectively, called “trainees”) with programs for professional development, career exploration, and career enhancement. Except for a few professional development sessions that are required for all first year PhD students, all other activities are optional and trainees choose from among the offerings according to their training stage and career interests. Professional development opportunities include a twice-monthly ASPIRE Postdoctoral Café series for postdocs, an annual ½ day ASPIRE to Connect workshop focused on the importance of building professional relationships, a series of career planning sessions for first year students or more advanced grad students and postdocs, and several non-credit bearing short courses in communication-related topics. Career exploration activities include a collection of nearly 100 Beyond the Lab interviews with Vanderbilt PhD and postdoctoral alumni discussing their careers (https://medschool.vanderbilt.edu/career-development/beyond-the-lab-see-listen/), a monthly PhD Career Stories seminar series, and an Annual Career Symposium. Career enhancement activities are intended for post-qualifying PhD students and postdoctoral fellows, and include opportunities to participate in didactic and experiential modules, the opportunity to gain hands-on-experience through ASPIRE internships, and the opportunity to gain deeper insight into specific industries through ASPIRE on the Road group field trips to visit cities with high concentrations of biotech or policy-related employers. For a description of the full range of ASPIRE program features, see the ASPIRE Annual Reports at https://medschool.vanderbilt.edu/career-development/annual-report/. |
| **Virginia Tech BEST Program:** | Virginia Tech’s Broadening Experiences in Scientific Training (BEST) Program activities are open to participation by pre- and postdoctoral scientists at any stage of their training, with some activities required by one or more graduate programs. Core offerings include: I. A 2-credit professional development course focused on self-assessment, skill-building, and career pathways for biomedical PhDs. Example skills topics include grantsmanship and CV writing, improvisation for science communication, and budgeting. II. Individual Development Planning workshops, III. Job simulation workshops delivered by outside professionals from a variety of careers, which involve hands-on activities and case studies. IV. A commercialization/shark tank module and pitch competition run jointly with biomedical engineering and business faculty. VT-BEST also delivers one-off activities such as networking training preceding professional/scientific events (with VT’s Career Center), and workshops such as visualizing data or social media for scientists (with VT’s Center for Communicating Science). Virginia Tech’s recent partnership with |
|  | the Roanoke RAMP accelerator provides training and shadowing opportunities in commercialization and start-ups. VT-BEST also facilitates a small number of internship opportunities, through industry partnerships and travel awards. Lastly, VT-BEST staff work closely with the Roanoke Graduate Student Association, Virginia Tech Carilion Student Outreach Program, and individual trainees, on the implementation of trainee-driven professional development activities. |
| **Wayne State BEST Program:** | Wayne State BEST assists doctoral students in exploring and pursuing a variety of career options. One of the highlights of Wayne State BEST is that doctoral students outside the biomedical disciplines also participate, adding richness to the training experiences. Wayne State BEST has three successive phases: Phase I – Exploratory Seminars; Phase II – Didactic Workshops; Phase III – Career Exploration/Internships. Phase I acquaints students with multiple career options via 90-minute seminars, each exploring one of five career tracks with industry partners, faculty, and alumni whose work intersects with the biosciences and the following areas: undergraduate teaching, law, communication, business/industry, and government. Phase II comprises a series of daylong workshops on the career options identified in Phase I. These workshops serve as a bridge between the Phase I exploratory seminars and the Phase III career exploration/internship experiences. A team comprising community and industry leaders (including alumni) work with faculty facilitators to design a curriculum focusing on necessary skills sets for each of the career tracks. Attendees gain additional knowledge through one-on-one exchanges with professionals in these domains. Students learn how their scientific training, problem-solving abilities, and analytical aptitude can be mobilized to successfully address the needs of their desired career. Phase III offers students experiential learning about these career paths through career explorations/internships with private industry, state agencies, nonprofit organizations, or primarily undergraduate institutions. Wayne State offers workshops on constructing an Individualized Development Plan (IDP), which is required of all doctoral students. In addition, the Wayne State Graduate School offers professional development seminars on basic employment skills such as conducting a job search, preparing for an interview, converting a CV to a resume, building a LinkedIn page, and writing a cover letter. Wayne State faculty lead specialized workshops on abstract writing, poster presentation, professional communication in the workplace, and strategies for presenting scientific ideas to non-specialist audiences. The Graduate School established a 1-credit course for graduate students interested in preparing for a career outside of academia. This course uses exercises and assignments to build a professional portfolio necessary for employment in highly skilled positions. |

### Supplemental File 2. Tables, and figures of participation effects on efficiency and productivity

**SI Table 2a.** Time to <u>degree</u> versus binary BEST participation (NTOTAL = 1700) In order to measure graduate students’ time in training, we contemplated if we should use time to degree or time to defense. One concern was that time to degree might not be a robust measure because it is a blunt instrument that can have delays built in between defending the dissertation and completing additional requirements, as well as delays due to official graduation dates. As a result, one would expect time to defense to be a more granular and sensitive measure to identify any potential delays due to involvement in professional development activities.

| **Institution** | **Mean**<br>**CTRL**<br>**Degree** | **Mean**<br>**BEST**<br>**Degree** | **R<sup>2</sup>** | **t-test** | **p-value** | **N**<br>**CTRL** | **N**<br>**BEST** | **N**<br>**TOTAL** |
| --- | --- | --- | --- | --- | --- | --- | --- | --- |
| Institution A | 73 | 69 | 0.01 | -1.78 | 0.08 | 101 | 136 | 237 |
| Institution B | 62 | 64 | 0.01 | +0.98 | 0.33 | 26 | 70 | 96 |
| **Institution C** | **73** | **68** | **0.04** | **-2.76** | **<0.01\*\*** | **50** | **126** | **176** |
| Institution D | 68 | 67 | <0.00 | -0.77 | 0.44 | 89 | 88 | 177 |
| Institution E | 69 | 69 | <0.00 | -0.09 | 0.93 | 121 | 102 | 223 |
| Institution F | 69 | 66 | <0.01 | -0.72 | 0.48 | 27 | 59 | 86 |
| Institution G | 52 | 56 | 0.02 | +1.24 | 0.22 | 79 | 20 | 99 |
| Institution H | 73 | 73 | <0.00 | -0.36 | 0.72 | 156 | 121 | 277 |
| Institution I | 69 | 68 | <0.00 | -0.76 | 0.45 | 106 | 132 | 238 |
| Institution J | 63 | 66 | <0.01 | 0.93 | 0.36 | 24 | 67 | 91 |

**SI Figure 2b.**
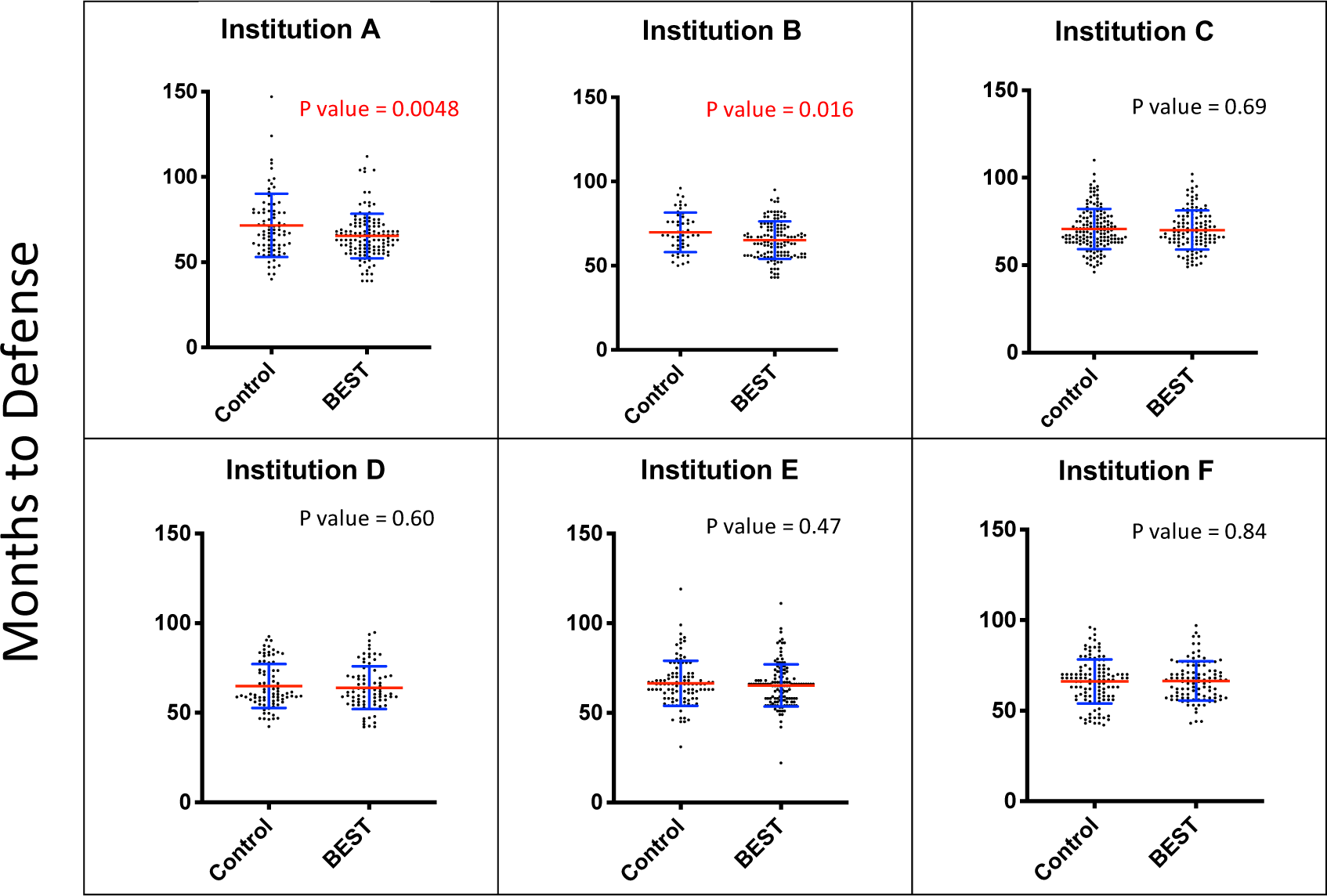
Time to <u>defense</u> versus binary professional development participation. Blue error bars represent standard deviation of the mean. Mean is denoted by a red line. Significant p-values (<0.05) are denoted in red whereas non-significant differences are denoted in black for each independent samples t-test.

**SI Table 2b.** Time to <u>defense</u> versus binary BEST participation (NTOTAL = 1300)

| Institution | M<br>CTRL<br>Defense | M<br>BEST<br>Defense | R<sup>2</sup> | t | p | N<br>CTRL | N<br>INT | N<br>TOT |
| --- | --- | --- | --- | --- | --- | --- | --- | --- |
| **Institution A** | **72** | **65** | **0.04** | **-2.85** | **<0.01** | **82** | **127** | **209** |
| **Institution B** | **70** | **65** | **0.03** | **-2.439** | **<0.02** | **50** | **126** | **176** |
| Institution C | 65 | 64 | <0.01 | -0.5266 | 0.60 | 89 | 88 | 177 |
| Institution D | 66 | 66 | <0.01 | +0.1980 | 0.84 | 121 | 102 | 223 |
| Institution E | 70 | 70 | <0.01 | -0.4042 | 0.69 | 156 | 121 | 277 |
| Institution F | 67 | 65 | <0.01 | -0.7178 | 0.47 | 106 | 132 | 238 |

**SI Figure 2c.**
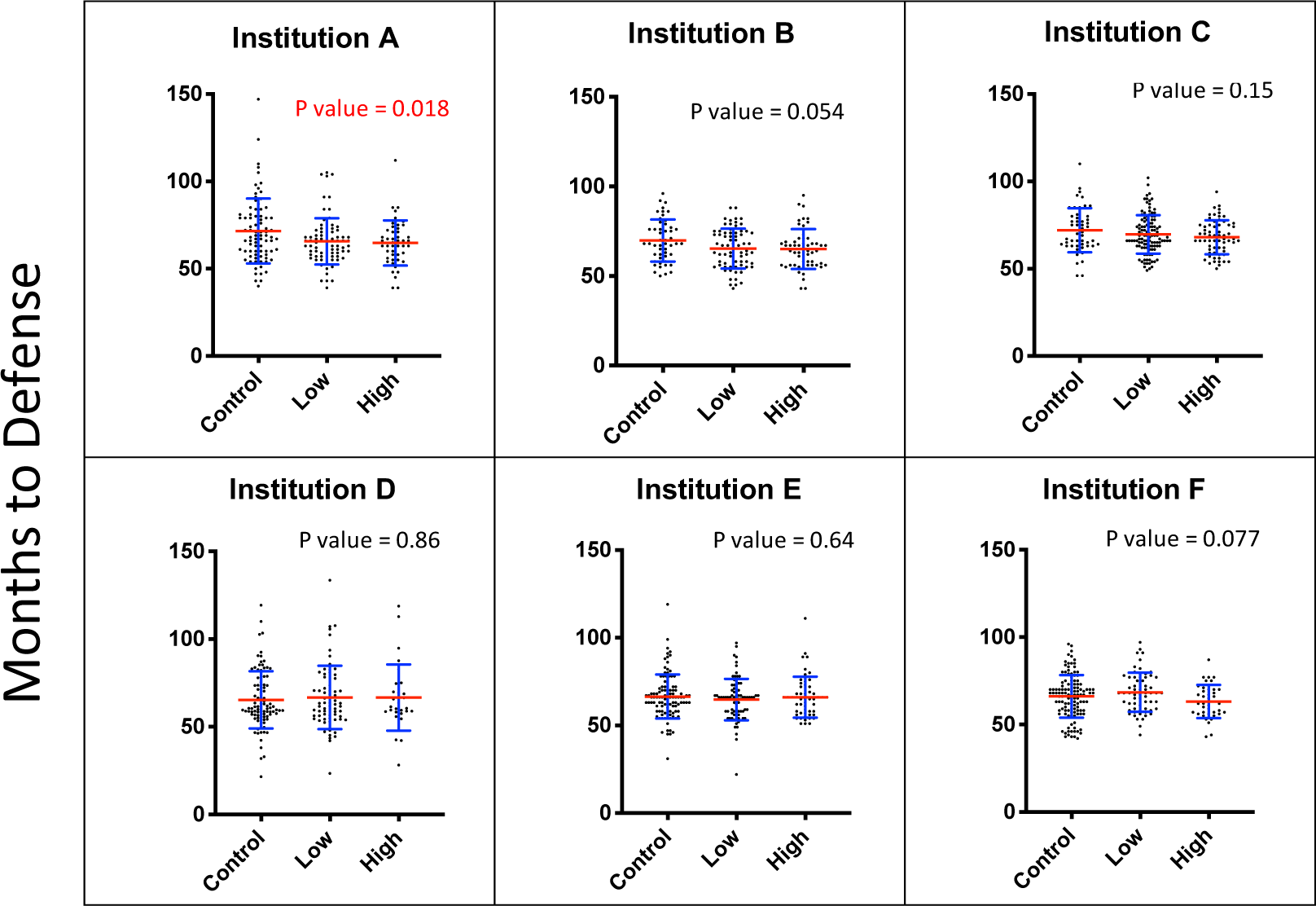
Time to <u>defense</u> versus dosage of professional development participation. Blue error bars represent standard deviation of the mean. Mean is denoted by a red line. Significant p-values (<0.05) are denoted in red whereas non-significant differences are denoted in black for each ANOVA (F-test).

**SI Table 2c.** Total Publications versus Professional Development Participation

| Institution | M<br>CTRL | M<br>BEST | R<sup>2</sup> | t | p-value | N<br>CTRL | N<br>BEST |
| --- | --- | --- | --- | --- | --- | --- | --- |
| **Institution A** | **3** | **4** | **0.02** | **+2.07** | **0.04\*** | **101** | **136** |
| Institution B | 4 | 4 | $<0.00$ | +0.25 | 0.80 | 26 | 71 |
| Institution C | 5 | 4 | 0.01 | -1.42 | 0.16 | 54 | 126 |
| **Institution D** | **6** | **4** | **0.02** | **-1.97** | **$<0.05^*$** | **97** | **97** |
| Institution E | 3 | 4 | $<0.00$ | -0.25 | 0.81 | 69 | 196 |
| Institution F | 6 | 5 | 0.01 | -1.04 | 0.30 | 25 | 52 |
| Institution G | 3 | 5 | 0.02 | +1.55 | 0.12 | 79 | 20 |
| Institution H | 5 | 5 | $<0.00$ | -1.00 | 0.32 | 52 | 197 |
| Institution I | 4 | 4 | $<0.01$ | -0.12 | 0.90 | 145 | 148 |

**SI Table 2d.** First-author Publications versus Professional Development Participation

**SI Table 2c.** Total Publications versus Professional Development Participation
| **Institution** | **M**<br>**CTRL**<br>**M<sub>CTRL</sub>**<br>**FA PUBLS** | **M**<br>**BEST**<br>**M<sub>BEST</sub>**<br>**FA PUBLS** | **R<sup>2</sup>** | **t** | **p-value** | **N**<br>**CTRL** | **N**<br>**BEST** |
| --- | --- | --- | --- | --- | --- | --- | --- |
| Institution A | 1 | 2 | <0.01 | +1.21 | 0.23 | 101 | 136 |
| Institution B | 2 | 1 | <0.01 | -0.56 | 0.57 | 26 | 71 |
| Institution C | 2 | 2 | <0.01 | -0.15 | 0.88 | 50 | 126 |
| **Institution D** | **2** | **2** | **0.03** | **-2.28** | **0.02\*** | **97** | **97** |
| Institution E | 1 | 1 | <0.01 | +1.58 | 0.12 | 69 | 207 |
| Institution F | 3 | 2 | 0.02 | -1.17 | 0.25 | 25 | 52 |
| Institution G | 2 | 2 | <0.00 | -0.30 | 0.76 | 79 | 20 |
| Institution H | 2 | 2 | <0.01 | -1.38 | 0.17 | 52 | 177 |
| Institution I | 2 | 2 | <0.01 | +0.05 | 0.96 | 145 | 148 |

**SI Table 2e.**
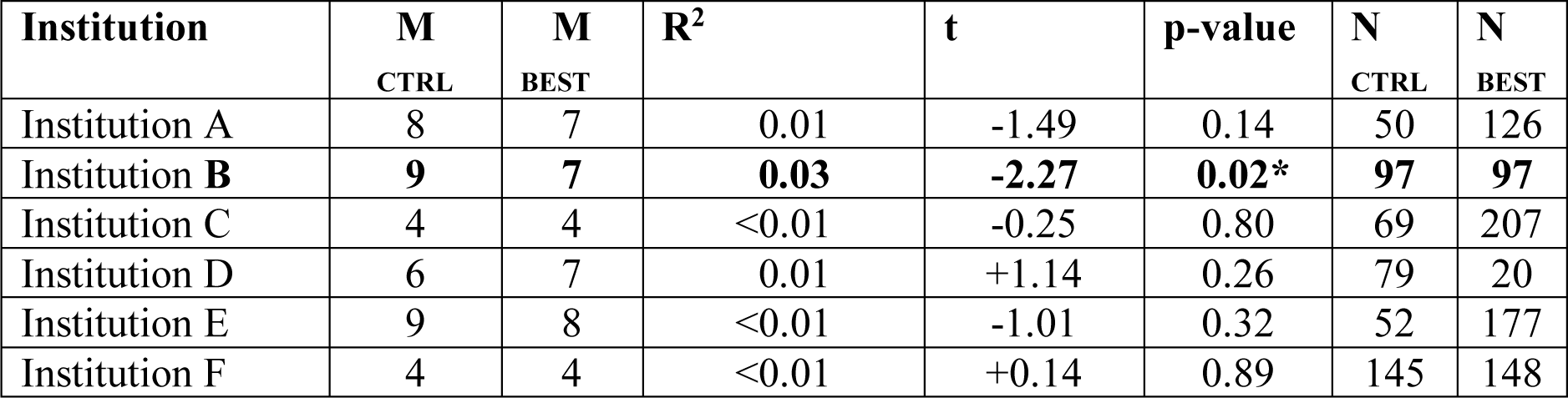
Publication Metric versus Professional Development Participation

### Supplemental Information - File 3. Figures and tables of internship effects on efficiency and productivity

**SI Table 3a.** Internship programs and definitions

| **Organization/**<br>**Institution** | **Experiential Learning Program Description** |
| --- | --- |
| ***NIH BEST Program:*** | Following the BEST Data Summit an internship is defined as working in a professional setting for the purpose of receiving *hands-on training*. An internship assumes the trainee is able to develop some skills during the experience and results in a deliverable. On the other hand, an externship is defined as job shadowing a professional at work for the purpose of *observing and experiencing* the work environment and learning about the expectations of a profession (O’Brien et al. in press). In both an internship and externship, significant time is spent in the professional workplace environment and therefore out of the graduate student’s own laboratory. For the purposes of this paper, internship and externship are collectively referred to as internship for consistency (exceptions are noted in figures as applicable). |
| ***Boston University BEST Program:*** | BU’s BEST program established relationships with departments and programs within BU and with local employers and nonprofit organizations to develop internship opportunities in diverse career tracks such as business, administration, communication, policy, research and teaching. Sites submitted a description of the internship including intern responsibilities, deliverable and its evaluation, professional development objectives, benefits to the intern, and assigned mentor. Internships were offered on a rolling basis, varied both in length (from 1 month to 1 year) and in time (from 1-40 hours/week), and were paid where possible. Applicants must have passed their qualifying exams prior to the start of an internship, have a completed IDP on file, and have attended basic skill building pre-internship workshops (e.g. professionalism 101) and internship-specific workshops. Applicants submitted their resume, approval by the research advisor, sign off on satisfactory academic performance, a personality assessment and a pre-internship evaluation. Each applicant met individually with the internship director prior and after applying to ensure that the trainee’s career goals were aligned with the goals of the internship. BU’s BEST presented qualified applicants to the internship site. Interns were selected by the employers, and developed projects with deliverables set by the |
|  | internship site. Both interns and site met with the internship director to evaluate learning objectives mid- and end of the internship. |
| ***Rutgers iJOBS Program:*** | Rutgers trainees who are interested in doing a deeper dive into a particular career track can apply to the Phase 2 cohort once they have completed at least 12 hours of Phase 1 events and completed their qualifying exams. About 20 trainees per year are admitted to Phase 2 and are matched by the iJOBS program directors with a professional in their area of interest for an externship/shadowing experience. The trainees spend time in the professional's workplace sitting in on meetings and observing their activities for a total of 72 hours spread out over the course of a semester. |
| ***University of Chicago myCHOICE Program:*** | A key goal of myCHOICE Experience programming is to provide real-world, practical experience in a specific career field. Internships, defined as "hands-on opportunities of limited duration (weeks to months)," are an important component of this training experience. myCHOICE collaborates with on- and off-campus partners to develop a diverse array of internships varying from scientific writing, to investment banking, to marketing and program development. All myCHOICE internships are unpaid and part-time (10 hours per week), lasting approximately 10 weeks. Trainees interested in internships must apply and receive permission from their PI. Interns who are graduate students formally register for the internship and receive academic credit. |
| ***University of North Carolina at Chapel Hill TIBBS Program:*** | UNC's ImPACT Internship program consisted of 160-hours, typically completed either one-month full time or multi-month part-time, in the career field of choice (industry research and development, science policy, teaching, museums/outreach, startups, etc.) Approximately 25-30 interns per year are typically selected to participate based on training stage (comps/quals complete to reach candidacy stage, often fourth or fifth years of training); research status (appropriate progress toward or completion of first-author publication for training stage); career exploration and professional development training; and selection of career path with competitive skills appropriate for field selected. This is the capstone experience available to UNC graduate students on a competitive basis are 160-hour internships that can take place during 1 month of full-time effort, or part time over 2-3 months. Interns are paid at their current stipend of salary rate. Graduate students must have passed their qualifying exams and all scientists must have written support of their faculty mentor in order to apply. |
| ***University of Rochester URBEST Program:*** | URBEST developed a flexible experiential learning program that included long-term internships (full-time, up to three month), short-term internships (hours-per-week) shadowing experiences (a couple of days total) and volunteer opportunities (< 4 hours per week). Internships took place within University of Rochester at core facilities (e.g., Office of Regulatory Support, Upstate Stem Cell Good Manufacturing Practice (GMP) Facility, Flow Cytometry Core) or within the city of Rochester (e.g., Rochester Museum and Science Center, Litron Laboratories). The majority of URBEST internships took place in different cities (e.g., Entasis Therapeutics, The US Food and Drug Administration (FDA), Pfizer Vaccines). To be allowed to do an internship, the learner must have officially enrolled in URBEST as a trainee for at least 6 months, have collected ~ 40 – 60 points through the program, have passed their qualifying exam and have the permission of their PI. The graduate student was also required to have a first author publication or multiple publications if they were not first author. The URBEST program disseminated intern opportunities as they became available, posting them first to our URBEST LinkedIn Group as a benefit of enrolling in the program. While undertaking their internship all trainees needed to be registered as a PhD graduate student. If the graduate student was on some type of training grant or fellowship, they needed to discuss your stipend and training opportunity with their program officer to get approval for their internship. It is up to the program officer as to whether or not their internship contributed to graduate student training, during the URBEST program all program officers approved graduate student internship requests. Most internships were set up by the trainee using a “cold email” technique to set up informational interviews, which often led to experiential opportunities. A few trainees found well established internships that they could also apply (e.g., Scientific American, Bayer Global Regulatory Affairs, White House Office of Science and Technology Policy). |
| ***Vanderbilt ASPIRE Program:*** | Vanderbilt University’s ASPIRE program has established relationships with local employers and several national nonprofit organizations to develop part-time internship opportunities in a range of career areas, including data science, college teaching, nonprofit management, business development, marketing, science policy and advocacy, and science outreach. ASPIRE internships are paid where possible, part-time (usually 6-8 hours per week), and generally last 10-12 weeks. Internships are offered on a rolling basis according to employer need and desired timing. All PhD student interns must have passed their qualifying exams prior to the start of an internship, and each applicant meets individually with our office staff prior to applying to ensure that the trainee’s career interests are aligned with the goals of the internship. Interns are selected by the employers, and interns are expected to contribute to one or more projects during their internship. Since the inception of the ASPIRE internship program in 2015, nearly 100 trainees have completed internships, about 65% of whom have been PhD students. |

**SI Table 3b.** Internships versus time to degree

| **Institution** | **M**<br>CTRL | **M**<br>INT | **R<sup>2</sup>** | **t-test** | **p-value** | **N**<br>CTRL | **N**<br>INT | **N**<br>TOTAL |
| --- | --- | --- | --- | --- | --- | --- | --- | --- |
| Institution A | 71 | 66 | <0.01 | -1.34 | 0.18 | 218 | 19 | 247 |
| Institution B | 70 | 66 | <0.01 | -1.57 | 0.12 | 155 | 21 | 176 |
| Institution C | 70 | 65 | 0.01 | -1.57 | 0.12 | 155 | 21 | 176 |
| Institution D | 72 | 76 | <0.00 | +1.09 | 0.28 | 237 | 12 | 249 |
| Institution E | 72 | 63 | <0.01 | -1.46 | 0.15 | 285 | 7 | 293 |
| Institution F | 69 | 67 | <0.01 | -0.82 | 0.41 | 196 | 28 | 223 |

**SI Table 3c.**
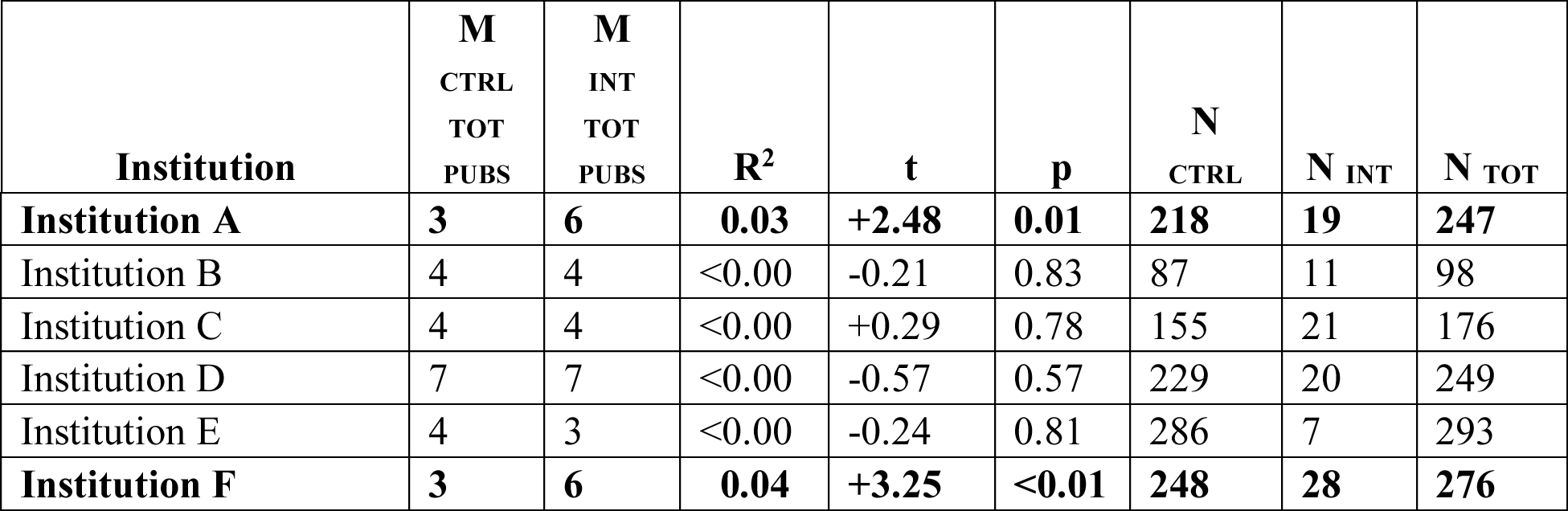
Internships versus total publications

**SI Table 3b. Internships versus time to degree**
| **Institution** | **M**<br>CTRL<br>TOT<br>PUBS | **M**<br>INT<br>TOT<br>PUBS | **R<sup>2</sup>** | **t** | **p** | **N**<br>CTRL | **N**<br>INT | **N**<br>TOT |
| --- | --- | --- | --- | --- | --- | --- | --- | --- |
| **Institution A** | **3** | **6** | **0.03** | **+2.48** | **0.01** | **218** | **19** | **247** |
| Institution B | 4 | 4 | <0.00 | -0.21 | 0.83 | 87 | 11 | 98 |
| Institution C | 4 | 4 | <0.00 | +0.29 | 0.78 | 155 | 21 | 176 |
| Institution D | 7 | 7 | <0.00 | -0.57 | 0.57 | 229 | 20 | 249 |
| Institution E | 4 | 3 | <0.00 | -0.24 | 0.81 | 286 | 7 | 293 |
| **Institution F** | **3** | **6** | **0.04** | **+3.25** | **<0.01** | **248** | **28** | **276** |

**SI Table 3d.** Internships versus first-author publications

**SI Table 3c. Internships versus total publications**
| **Institution** | **M**<br>**CTRL**<br>**FA**<br>**PUBS** | **M**<br>**INT FA**<br>**PUBS** | **R<sup>2</sup>** | **t** | **p** | **N**<br>**CTRL** | **N**<br>**INT** | **N**<br>**TOT** |
| --- | --- | --- | --- | --- | --- | --- | --- | --- |
| Institution A | 1 | 2 | <0.00 | +0.12 | 0.90 | 218 | 19 | 247 |
| Institution B | 4 | 2 | <0.04 | -2.03 | 0.05 | 86 | 11 | 97 |
| **Institution C** | **2** | **2** | **0.03** | **-2.32** | **0.02\*** | **155** | **21** | **176** |
| Institution D | 2 | 2 | <0.00 | +0.17 | 0.86 | 229 | 20 | 249 |
| Institution E | 2 | 2 | 0.01 | -0.56 | 0.58 | 286 | 7 | 293 |
| **Institution F** | **1** | **2** | **0.05** | **+3.71** | **<0.01** | **248** | **28** | **276** |

**SI Table 3e.** Internships versus publication metric

**SI Table 3d. Internships versus first-author publications**
| **Institution** | **M**<br>**CTRL**<br>**PUB MET** | **M**<br>**INT**<br>**PUB MET** | **R<sup>2</sup>** | **t** | **p** | **N**<br>**CTRL** | **N**<br>**INT** | **N**<br>**TOT** |
| --- | --- | --- | --- | --- | --- | --- | --- | --- |
| Institution A | 7 | 7 | <0.00 | +0.37 | 0.71 | 155 | 21 | 176 |
| Institution B | 8 | 7 | <0.00 | -0.71 | 0.48 | 229 | 20 | 249 |
| Institution C | 4 | 3 | 0.01 | -0.56 | 0.57 | 286 | 7 | 293 |
| **Institution D** | **4** | **6** | **0.04** | **+3.08** | **<0.01** | **248** | **28** | **276** |

**SI Figure 3a.**
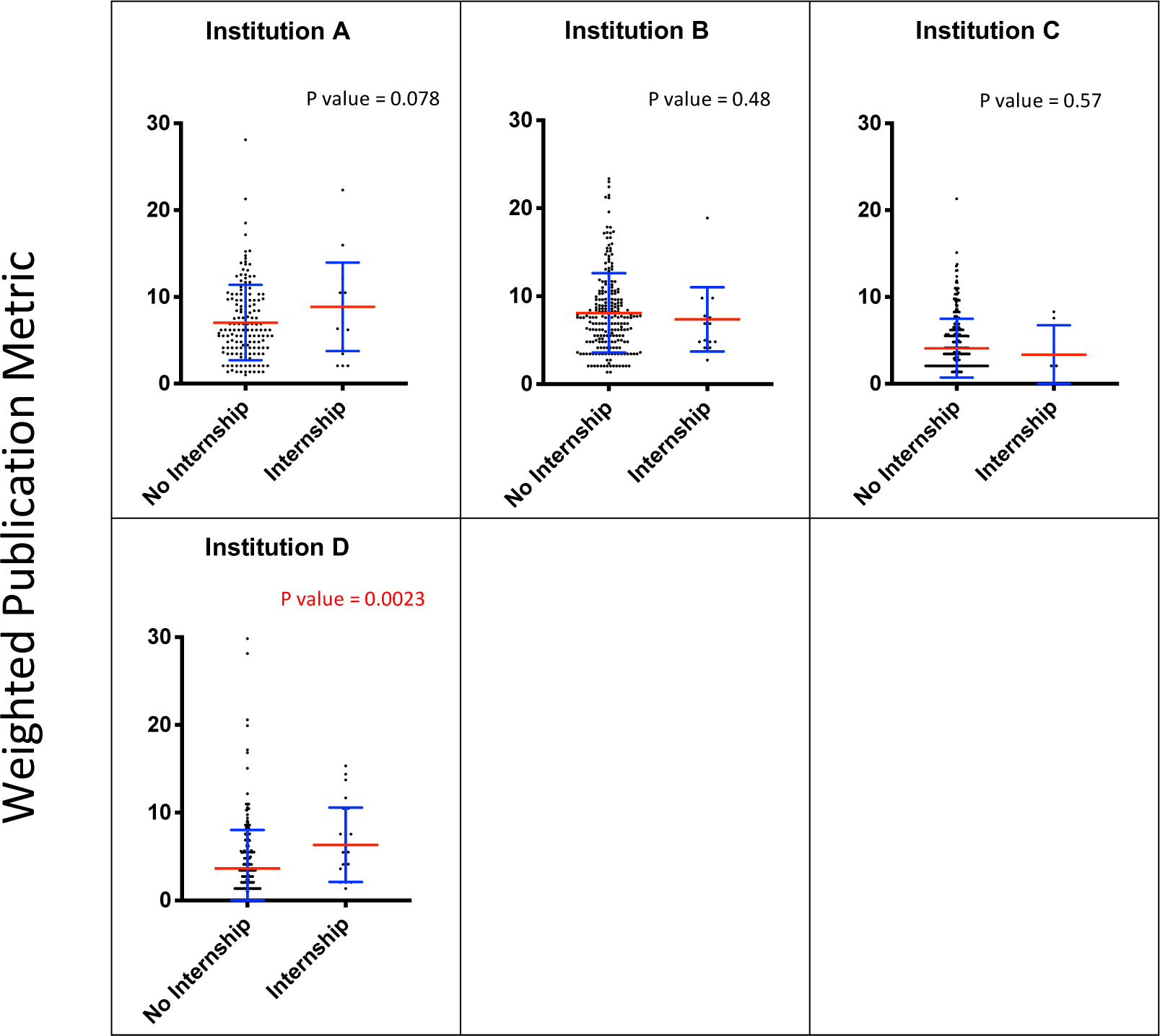
New publication metric versus internship/externship participation (both included in internship category herein). Blue error bars represent standard deviation of the mean. Mean is denoted by a red line. Non-significant differences are denoted in black for each independent samples t-test.

### Supplemental File 4. Publication reporting and metric development

#### Publication data collection procedures

First-author and co-first-author publications were included in the first-author publication count. Publications and their metadata were primarily collected through a Python script built to query the PubMed API (47), in combination with manual verification (see **SI Table 4a**). By automating the PubMed search process, the script allowed for replication and validation of publication data across multiple institutions and implementation of Cross-Institutional Instructions. Manual checking was used for institutions that could not access PubMetric results for technical reasons (one of ten institutions); or existing survey data from a graduate school survey were used (one of ten institutions).

#### Publication metric

The metric is a weighted calculation of different types of publications determined by polling 375 active training faculty at UNC about the relative value they place on different publication types. Respondents (n=120) were asked to assign their values to the following types of publications:

- First-author (and co-first-author) research paper (FA Res)
- Middle-author research paper (MA Res)
- First-author review (FA Rev)
- Middle-author review (MA Rev)

Based on the responses on a scale of 1-10 (1=*Not valuable*, 10=*Extremely valuable*), an average rating was calculated for each type of publication. Once all four of the most common metrics were calculated, a weighted publication metric was created to represent the productivity of any given trainee. The metric is given by the equation: 2.07*Number of First-author Research papers + 1.37*Number of Middle-author Research papers + 1.54*Number of First-author Reviews +1*Number of Middle-author Review papers.

The new weighted pub metric developed based on the average of 120 responses was as follows: = 2.07*(FA Res)+1.37*(MA Res)+1.54*(FA Rev)+1*(MA Rev)

This allows for the use of a single publication metric rather than having to depend on multiple measures, and may be of use especially when simplicity or overall trends are of most interest. This method of evaluating productivity is an alternative to attempting to assign credit to various flagship journals by name (which can be difficult to capture across fields), or impact factor measures (which are controversial), and provides an independent estimate of productivity based on role/contribution to each work as well as accounting for type of publication.

Sample survey

SURVEY: Faculty survey to create publication productivity rating

Start of Block: Productivity Metric Survey

As part of a project to examine graduate student productivity we need your input. Your response will be used to develop a new metric to represent trainee publication “productivity” as a single quantitative measure.

Please rate the relative value you give to each publication type when evaluating trainee productivity.

Q: How would you value the contribution of a candidate with each of the following publication types?

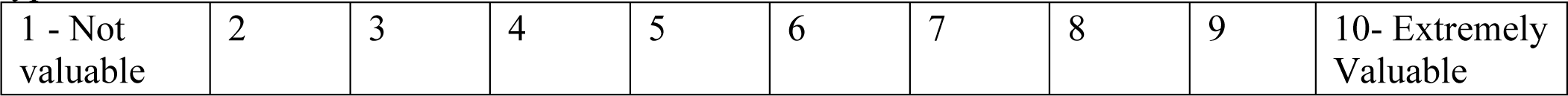

First-Author Peer-Reviewed Scientific Publication Middle-Author Peer-Reviewed Scientific Publication

First-Author Review Article or Book Chapter (Peer Reviewed) Middle-Author Review Article or Book Chapter (Peer Reviewed) Other publication type?(specify or skip)

Other publication type? (specify or skip) Other publication type? (specify or skip)

Q: Have you supervised graduate or post-doctoral trainees? (Yes/No)

Q: How many total graduate student and post-doctoral trainees have you supervised? Number of trainees (total)

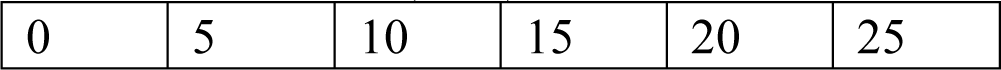

Thank you for helping to develop a new metric for assessing graduate student productivity. We will share the survey results with all respondents after the survey closes on 12/6/17.

**SI Table 4a.** PubMed Crawler Script – Data Integrity Measures by Institution

| **Institution** | **Publication Data Collection Method** | **Data check method** | **Note** |
| --- | --- | --- | --- |
| Institution A | Algorithm | Manual checks | Checked for 0 & > 4 |
| Institution B | Algorithm | Manual checks | Checked for 0 & > 4 |
| Institution C | Algorithm | Checked all | Checked all |
| Institution D | Algorithm | Checked all | Checked all |
| Institution E | Manual Lookup | Checked all | Checked all |
| Institution F | Surveys | Self-report | Student exit survey |
| Institution G | Algorithm | Checked all | Checked all |
| Institution H | Internal Algorithm | Manual checks | Checked for 0 and $\geq 15$<br>Common names verified |
| Institution I | Algorithm + Internal Algorithm | Manual checks | Checked for 0 & > 20 |
|  |  |  | 90 algorithm<br>120 internal algorithm |

**SI Figure 4a.**
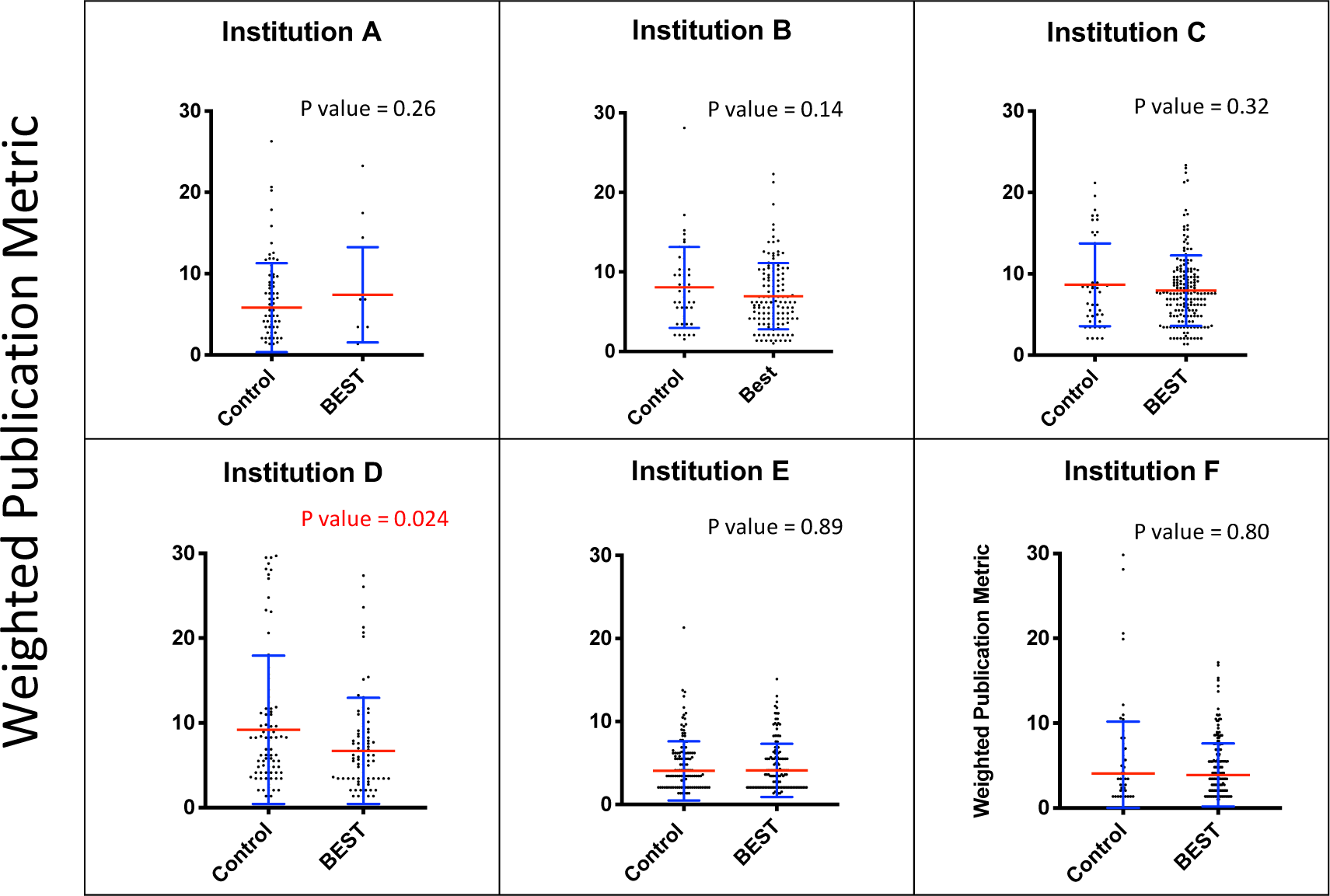
Weighted publication metric versus binary professional development participation. Blue error bars represent standard deviation of the mean. Mean is denoted by a red line. Significant p-values (<0.05) are denoted in red whereas non-significant differences are denoted in black for each independent samples t-test.

**SI Figure 4b.**
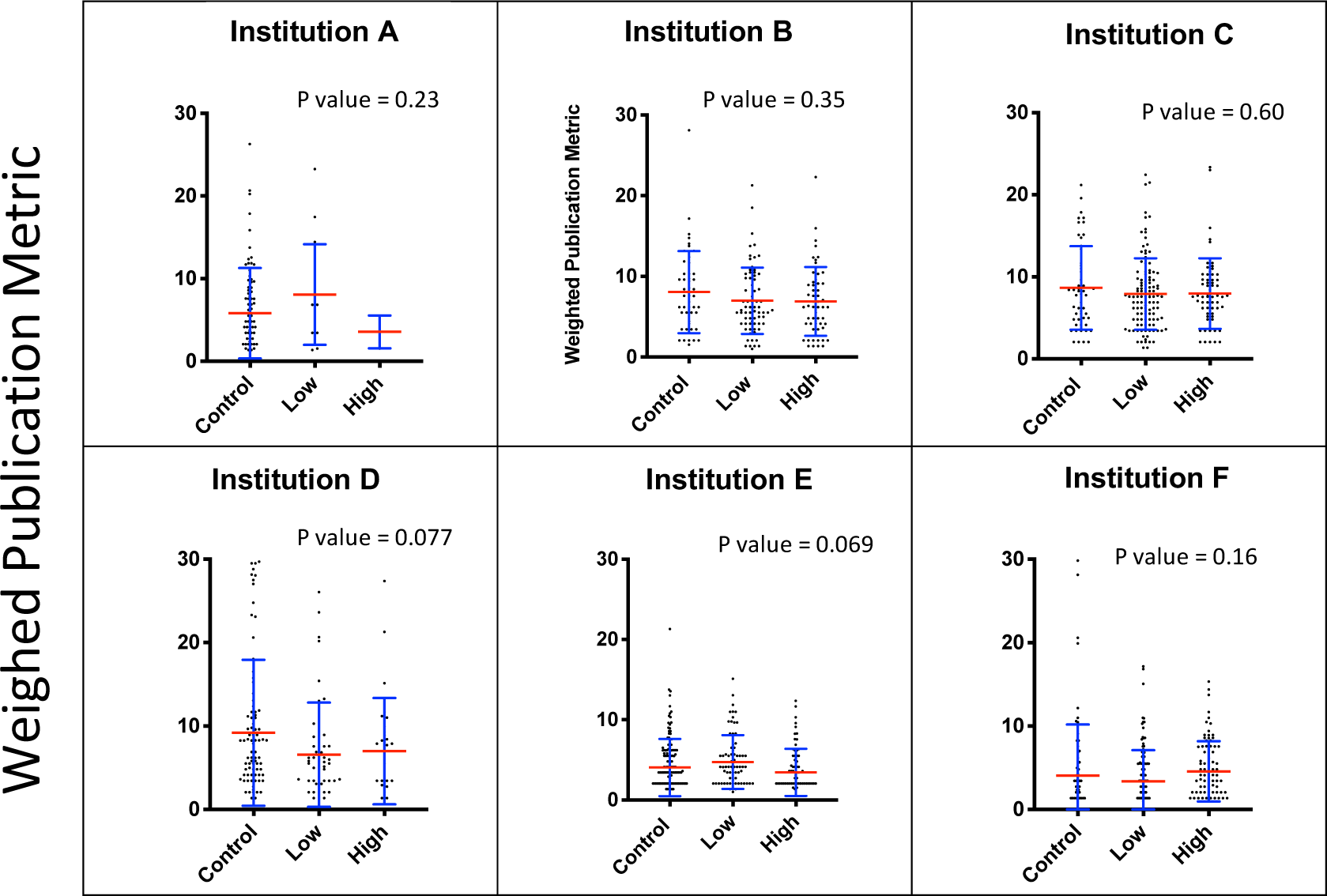
Weighted publication metric versus dosage of professional development participation. Blue error bars represent standard deviation of the mean. Mean is denoted by a red line. Non-significant differences are denoted in black for each ANOVA (F-test).

### Supplemental File 5. Comparison data for rotations

**SI Table 5.**
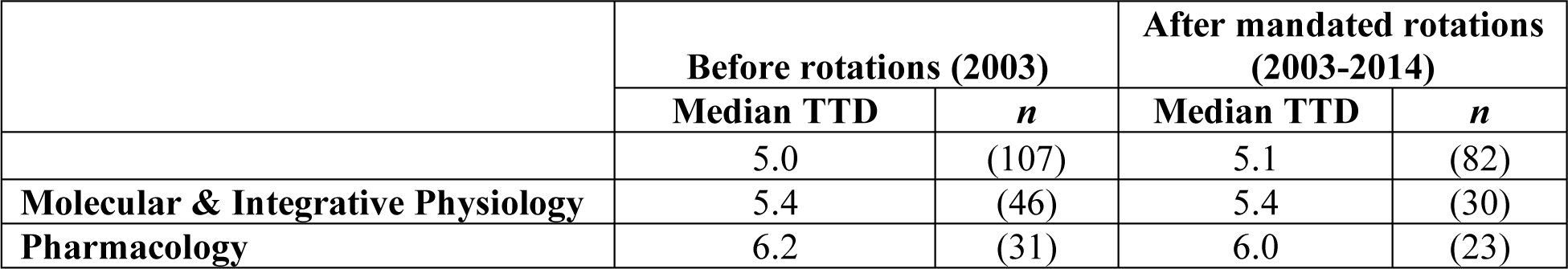
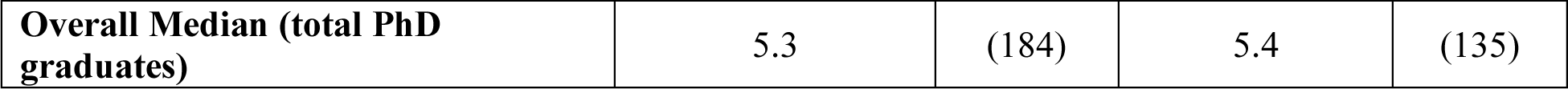
**Time to degree (in years) versus rotations** Data analyzed from Cornell University revealed no statistically significant lengthening of degree across comparison groups before and after rotations were mandated in 2003 for three graduate fields.SI Table 5 **Legend.** Independent samples t-tests (*t* = 0.80, *p* = NS) do not show a significant difference (raw data available in Supplemental Information on the <u>Open Science Framework repository</u>).

### Supplemental File 6. Data

Supplemental Information – Data Sets: SPSS Files

Available: <u>Open Science Framework repository</u>

## References

1. B. Bryan, K. Guccione, Was it worth it? A qualitative exploration into graduate perceptions of doctoral value. High. Educ. Res. Dev. 37, 1124–1140 (2018).

2. K. A. Petrie, R. H. Carnahan, A. M. Brown, K. L. Gould, Providing Experiential Business and Management Training for Biomedical Research Trainees. CBE—Life Sci. Educ. 16, ar51 (2017).

3. 3. NIH, NIH Biomedical Workforce Working Group Report. Available at https://acd.od.nih.gov/documents/reports/Biomedical_research_wgreport.pdf. (2012).

4. B. Alberts, M. W. Kirschner, S. Tilghman, H. Varmus, Rescuing US biomedical research from its systemic flaws. Proc. Natl. Acad. Sci. 111, 5773–5777 (2014).

5. NSF, Doctorate Recipients from U.S. Universities | NCSES | NSF. Available at https://www.nsf.gov/statistics/doctorates/. (July 10, 2020).

6. K. LanginMar. 12, 2019, 5:45 Pm, In a first, U.S. private sector employs nearly as many Ph.D.s as schools do. Sci. AAAS (2019) (July 10, 2020).

7. C. N. Fuhrmann, Enhancing Graduate and Postdoctoral Education To Create a Sustainable Biomedical Workforce. Hum. Gene Ther. 27, 871–879 (2016).

8. R. N. Lenzi, S. J. Korn, M. Wallace, N. L. Desmond, P. A. Labosky, The NIH “BEST” programs: Institutional programs, the program evaluation, and early data. FASEB J. 34, 3570–3582 (2020).

9. A. I. Leshner, Student-centered, modernized graduate STEM education. Science 360, 969– 970 (2018).

10. M. J. Mulvany, Biomedical PhD Education - An International Perspective. Basic Clin. Pharmacol. Toxicol. 112, 289–295 (2013).

11. NSF, National Science Foundation Research Traineeship (NRT) Program | NSF - National Science Foundation. Available at https://www.nsf.gov/funding/pgm_summ.jsp?pims_id=505015. (July 10, 2020).

12. J. Lorsch, A. Gammie, S. Singh, L. Newman, L. Moeller, NIGMS Ruth L. Kirschstein NRSA Predoctoral Institutional Research Training Grant (T32). 64 (2018).

13. NIGMS, NIGMS Predoctoral Training Grant Program Areas and Contacts. Available at https://www.nigms.nih.gov/training/instpredoc/pages/PredocDesc-Contacts.aspx. (July 21, 2020).

14. E. National Academies of Sciences, The Next Generation of Biomedical and Behavioral Sciences Researchers: Breaking Through (2018) (July 10, 2020).

15. D. Chatterjee, J. K. Ford, J. Rojewski, S. W. Watts, Exploring the impact of formal internships on biomedical graduate and postgraduate careers: An interview study. CBE Life Sci. Educ. 18 (2019).

16. J. M. Colby, F. C. Wheeler, K. A. Petrie, K. L. Gould, J. E. Schmitz, Institutional Training Opportunities for PhD Students in Laboratory Medicine: An Unmet Career Development Need? J. Appl. Lab. Med. 5, 412–416 (2020).

17. AAAS, Homepage - AAAS. Available at https://careerdevelopment.aaas.org/. *AAAS Career Dev. Cent*. (July 10, 2020).

18. AAAS, Catalyzing Advocacy in Science and Engineering Workshop | American Association for the Advancement of Science. Available at https://www.aaas.org/programs/catalyzing-advocacy-in-science-and-engineering. (July 21, 2020).

19. ASCB, Career Development ASCB. Available at https://www.ascb.org/career-development/. ASCB (July 21, 2020).

20. Genetics Society of America, Career Development Genetics Society of America. Available at https://genetics-gsa.org/career-development/. Genet. Soc. Am. (July 21, 2020).

21. L. Gillingham, J. J. Seneca, M. K. Taussig, The determinants of progress to the doctoral degree. Res. High. Educ. 32, 449–468 (1991).

22. F. J. Elgar, R. M. Klein, What You Don’t Know: Graduate Deans’ Knowledge of Doctoral Completion Rates. High. Educ. Policy 17, 325–336 (2004).

23. P. Hitchcock, et al., The future of graduate and postdoctoral training in the biosciences. eLife 6, e32715 (2017).

24. S. Kahn, D. K. Ginther, The impact of postdoctoral training on early careers in biomedicine. Nat. Biotechnol. 35, 5 (2017).

25. S. W. Watts, et al., Faculty perceptions and knowledge of career development of trainees in biomedical science: What do we (think we) know? PLOS ONE 14, e0210189 (2019).

26. L. Ullrich, et al., From student to steward: the Interdisciplinary Program in Neuroscience at Georgetown University as a case study in professional development during doctoral training. Med. Educ. Online 19, 22623 (2014).

27. D. B. Noble, S. G. J. Mochrie, C. S. O’Hern, T. D. Pollard, L. Regan, Promoting convergence: The integrated graduate program in physical and engineering biology at Yale University, a new model for graduate education: Promoting Convergence. Biochem. Mol. Biol. Educ. 44, 537–549 (2016).

28. D. F. Feldon, et al., Graduate Students’ Teaching Experiences Improve Their Methodological Research Skills. Science 333, 1037–1039 (2011).

29. E. E. Shortlidge, S. L. Eddy, The trade-off between graduate student research and teaching: A myth? PLOS ONE 13, e0199576 (2018).

30. M. Roach, Encouraging entrepreneurship in university labs: Research activities, research outputs, and early doctorate careers. PLOS ONE 12, e0170444 (2017).

31. A. Mathur, F. J. Meyers, R. Chalkley, T. C. O’Brien, C. N. Fuhrmann, Transforming training to reflect the workforce. Sci Transl Med 7, 285ed4 (2015).

32. C. J. van Lissa, MetaForest: Exploring heterogeneity in meta-analysis using random forests (2017) https:/doi.org/10.31234/osf.io/myg6s (July 10, 2020).

33. J. Cohen, Statistical power analysis for the behavioral sciences, 2nd ed (L. Erlbaum Associates, 1988).

34. J. Cohen, Statistical Power Analysis. Curr. Dir. Psychol. Sci. 1, 98–101 (1992).

35. M. Harrer, P. Cuijpers, T. A. Furukawa, D. D. Ebert, Doing Meta-Analysis in R (July 9, 2020).

36. M. Borenstein, L. V. Hedges, J. P. T. Higgins, H. R. Rothstein, Introduction to Meta-Analysis | Wiley. Wiley.com (2011) (July 9, 2020).

37. J. D. Hall, A. B. O’Connell, J. G. Cook, Predictors of Student Productivity in Biomedical Graduate School Applications. PLOS ONE 12, e0169121 (2017).

38. L. Moneta-Koehler, A. M. Brown, K. A. Petrie, B. J. Evans, R. Chalkley, The Limitations of the GRE in Predicting Success in Biomedical Graduate School. PLOS ONE 12, e0166742 (2017).

39. V. Mangematin, PhD job market: professional trajectories and incentives during the PhD. Res. Policy 29, 741–756 (2000).

40. A. M. Schnoes, et al., Internship Experiences Contribute to Confident Career Decision Making for Doctoral Students in the Life Sciences. CBE Life Sci. Educ. 17, 1–14 (2018).

41. NIH, NOT-OD-15-008: OMB Clarifies Guidance on the Dual Role of Student and Postdoctoral Researchers. Available at https://grants.nih.gov/grants/guide/notice-files/NOT-OD-15-008.html. (July 10, 2020).

42. F. J. Meyers, et al., The origin and implementation of the Broadening Experiences in Scientific Training programs: An NIH common fund initiative. FASEB J. 30, 507–514 (2016).

43. R. S. Clair, et al., The “new normal”: Adapting doctoral trainee career preparation for broad career paths in science. PLOS ONE 12, e0177035 (2017).

44. S. Gehr, C. C. Garner, K. N. Kleinhans, Translating academic careers into industry healthcare professions. Nat. Biotechnol. 38, 758–763 (2020).

45. Burroughs Wellcome Fund, Career Guidance for Trainees | Burroughs Wellcome Fund. Available at https://www.bwfund.org/grant-programs/career-guidance/career-guidance-trainees. (July 21, 2020).

46. L. I. Lara, L. Daniel, R. Chalkley, BEST: Implementing Career Development Activities for Biomedical Research Trainees, 1st Ed. (2020).

47. J.D. Hall & D. Arneman. PubMed authorship crawler. GitHub, available: https://github.com/dkarneman/pubmetric (2020).

48. H. Singh & D.A. Fruman. “Keys to successful implementation of a professional development program: Insights from UC Irvine’s GPS-BIOMED.” In BEST, pp. 129–137. Academic Press (2020).

49. A. Van Wart, T.C. O’Brien, S.S. Varvayanis, J. Alder, J. Greenier, R.L. Layton, C.A. Stayart, I. Wefes, A.E. Brady (in press).. Applying Experiential Learning to Career Development Training for Biomedical Graduate Students and Postdocs: Perspectives on Program Development and Design. CBE—Life Sciences Education.

